# Spatiotemporal analysis of dystrophin expression during muscle repair

**DOI:** 10.1101/2024.12.06.627177

**Authors:** John C.W. Hildyard, Liberty E. Roskrow, Dominic J. Wells, Richard J. Piercy

## Abstract

Dystrophin mRNA is produced from a very large genetic locus and transcription of a single mRNA requires approximately 16 hours. This prolonged interval between transcriptional initiation and completion results in unusual transcriptional behaviour: in skeletal muscle, myonuclei express dystrophin continuously and robustly, yet degrade mature transcripts shortly after completion, such that most dystrophin mRNA is nascent, not mature. This implies dystrophin expression is principally controlled post-transcriptionally, a mechanism that circumvents transcriptional delay, allowing rapid responses to change in demand. Dystrophin protein is however highly stable, with slow turnover: in healthy muscle, despite constant production of dystrophin mRNA, demand is low and the need for responsive expression is minimal. We reasoned this system instead exists to control dystrophin expression during rare periods of elevated but changing demand, such as during muscle development or repair, when newly formed fibres must establish sarcolemmal dystrophin rapidly. By assessing dystrophin mRNA and protein expression in regenerating skeletal muscle following injury, we reveal a complex program that suggests control at multiple levels: nascent transcription begins even prior to myoblast fusion, effectively ‘paying in advance’ to minimise subsequent delay. During myotube differentiation and maturation, when sarcolemmal demands are high, initiation increases only modestly while mature transcript stability increases markedly to generate high numbers of mature dystrophin transcripts, a state that persists until repair is complete, when a state of oversupply and degradation resumes. Our data demonstrate that dystrophin mRNA is indeed chiefly controlled by turnover, not initiation: degradation consequently represents a potential therapeutic target for maximising efficacy of even modest dystrophin restoration.

## Introduction

Dystrophin is essential for muscle health: in skeletal muscle myofibres this 427kDa protein (dp427) is found at the sarcolemma as a core component of the dystrophin associated glycoprotein complex (DAGC), a macromolecular assembly that links cytoskeletal actin and microtubule networks to the extracellular matrix (ECM). The complex forms a physical bridge between the myofibre cytoskeleton and the ECM environment, distributing contractile force and buffering the sarcolemmal membrane against the stress of muscle activity (Figure 1A). Without dp427, myofibres are vulnerable to contraction-induced damage, and loss or insufficiency of this protein results in the severe muscle wasting condition Duchenne muscular dystrophy (DMD), a fatal, incurable disease characterised by repeated cycles of muscle damage, regeneration and concomitant inflammation, fibrosis and atrophy.

**Figure 1:**
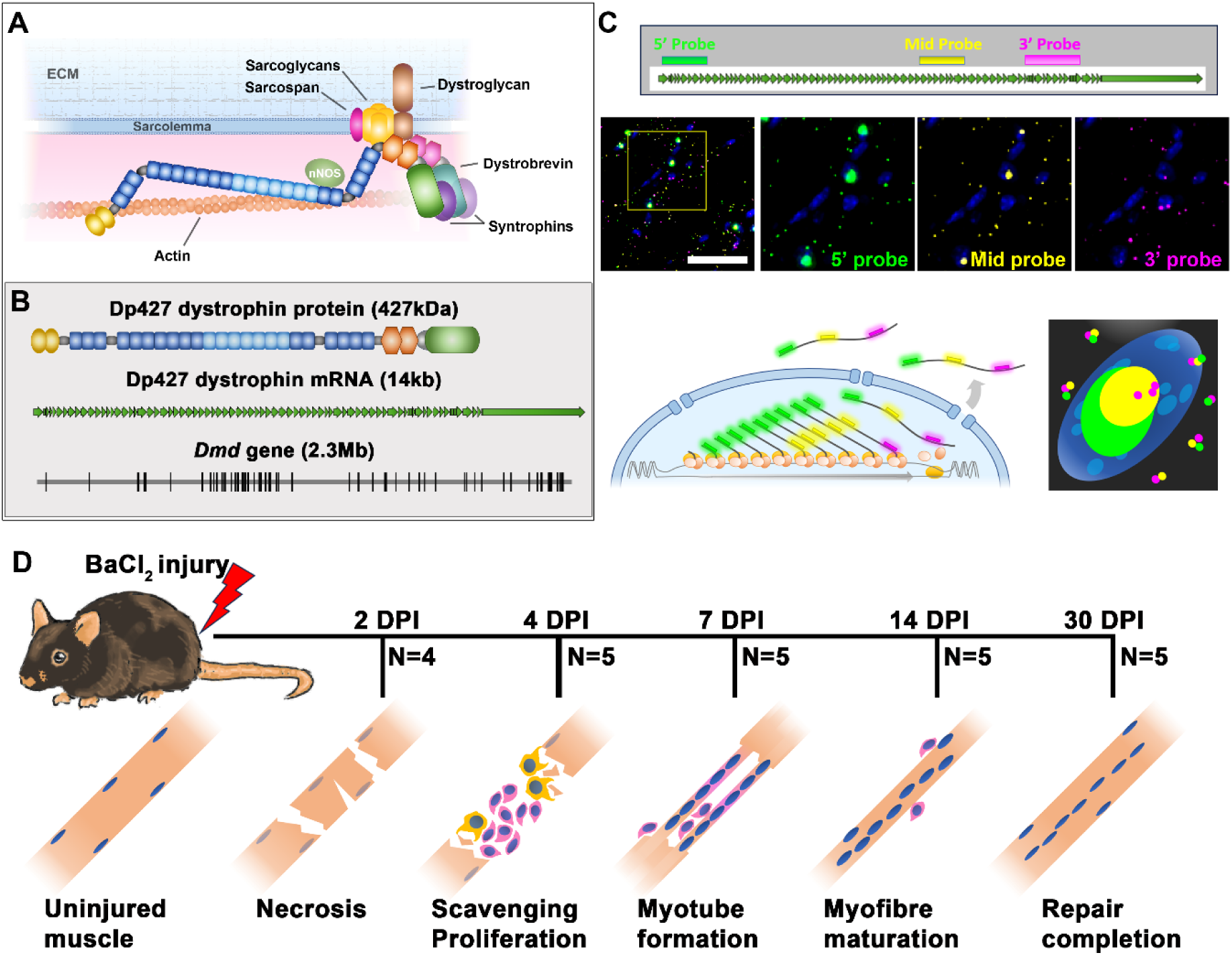
Overview. (A) Dystrophin (dp427) protein localises to the sarcolemma as a core component of the dystrophin-associated glycoprotein complex (DAGC), physically connecting cytoskeletal actin to sarcolemmal-spanning dystroglycan, and localising multiple additional proteins. (B) Sizes/lengths of dystrophin sequences: dp427 protein (∼1300 amino acids, 427kDa) and mRNA (14kb) are very large, but the Dmd gene (2.3Mb) is enormous. (C) Use of multiplex RNAscope FISH probes to dystrophin 5’, middle and 3’ regions reveals a unique labelling pattern consistent with large numbers of nascent transcripts within myonuclei, while mature transcript numbers are more modest (Scalebar: 50µm). Experimental model (D): WT mice were injured via BaCl_2_ injection (left tibialis anterior), and muscles (injured and uninjured) were collected at the indicated days post injury (DPI), consistent with necrosis (day 2), proliferation/scavenging (day 4), early regeneration (day 7), late regeneration (day 14) and repair completion (day 30). Animal numbers for each time-point are indicated.

Dystrophin is transcribed from an exceptionally long gene (*Dmd*): at ∼2.3 megabases, it is one of the longest genes in the mammalian genome, a feature that presents significant biological challenges (Figure 1B). Much of the gene is intronic (several introns are >100kb) and the 14kb mature mRNA represents less than 0.5% of total *Dmd* sequence. However, production of a single dp427 transcript still requires all 2.3Mb of the *Dmd* gene to be transcribed in full, without RNA polymerase dissociation. Work by Tennyson *et al.* in the 1990s demonstrated that transcription of a single dystrophin mRNA requires approximately 16 hours [1], implying a mean transcription rate of 40 bases.sec^-1^ (a value that accords well with estimates for other long genes [2, 3]). This lengthy transcription time therefore likely represents essentially uninterrupted polymerase activity, and indeed work by Gherardi *et al* [4] confirmed that polymerase pausing along the *Dmd* locus was not a significant contributor to this long transcription time. Tennyson *et al.* further showed that splicing of dp427 occurs co-transcriptionally, with mature, spliced 5’ sequence detectable significantly before the appearance of any 3’ sequence [1], and work by Gazzoli *et al.* demonstrated this splicing is complex [5], with specific groups of exons spliced before others (with some particularly long introns even spliced out in sequential, piecemeal fashion). A remarkable conclusion of these early studies (and also reported by subsequent investigations) is that 5’ sequence is not simply transcribed several hours before 3’ sequence, but also remains in excess of 3’ sequence persistently: indeed, in mature, healthy skeletal muscle where dystrophin expression should be essentially at steady-state, spliced dystrophin 5’ sequence (exons 1-2) can reliably be detected at markedly higher levels than 3’ (exons 62-63) [6, 7]. This phenomenon has been observed with dystrophin in humans, mice and dogs, and has been termed ‘transcript imbalance’ [8]. Imbalance is typically more dramatic in dystrophic muscle, leading to further suggestions that it might be pathologically relevant. An alternative hypothesis is that this reflects inherent instability of mature dp427 mRNAs, with a half-life of some ∼4 hours, substantially shorter than the 16 hours required for production. Under such a scenario, initiated but not-yet-complete dp427 transcripts (bearing spliced 5’ sequence but not 3’) might be present in considerable numbers, while mature transcripts (bearing both spliced 5’ and 3’ sequence) would be in the minority. In essence, even under steady-state healthy conditions, most dp427 mRNA would be nascent, not mature.

We have previously demonstrated that this latter scenario is indeed the case, by using a multiplex single-molecule fluorescence *in situ* hybridisation (smFISH) approach to label different regions (5’, central and 3’) of the long dp427 mRNA [7, 9]. Within healthy mature (multinucleated) skeletal muscle fibres, individual mature sarcoplasmic transcripts are present in modest numbers (labelled in a similar punctate manner with all three probes) and are predominantly localised to the sarcolemma. Myonuclei, in contrast, are host to large, intense foci of dystrophin 5’ probe, but only scattered punctate foci of 3’ probe, while signal from probes to the central dp427 sequence falls between these two extremes (Figure 1C). This labelling pattern is consistent with high numbers of immature mRNAs (∼20-40 per nucleus [7]) at various stages of transcription. Most dp427 mRNA is indeed nascent, not mature, and ‘transcript imbalance’ appears to be simply an inevitable consequence of robust and ongoing transcriptional initiation, long transcription time, and short mature mRNA half-life (discussed in more detail elsewhere [10]).

This transcriptional model is puzzling, implying near-continuous initiation of dystrophin transcription, ongoing commitment to the 16 hours (and 2.3 million base incorporation events) required for each transcript, yet also comparatively prompt degradation of those transcripts once completed. Added to this, the turnover of dystrophin protein in skeletal muscle is very slow (weeks to months [11, 12]): the majority of these continuously produced transcripts might thus be wholly surplus to requirements. Supply of dystrophin mRNA appears to substantially outstrip demand, and in healthy muscle at steady state, it seems plausible that the bulk of dp427 mRNA produced is not meaningfully translated at all (this translational inactivity could itself potentiate transcript degradation [13, 14]).

We proposed [10] that this system allows fine tuning of dystrophin transcripts over short timescales: instead of initiating transcription in response to demand (a process that would necessarily incur a 16 hour delay), myonuclei express dystrophin constitutively, regardless of demand, and mRNA levels are instead adjusted post-transcriptionally via control of degradation. By sacrificing efficiency for control, increases in demand could be met almost immediately by simply stabilizing newly completed transcripts that were initiated 16 hours previously. Under this model, myonuclei would effectively ‘pay in advance’ continuously under the assumption that dystrophin mRNA is always needed at high levels, and then degrade that mRNA subsequently should it transpire that demand is low. While this represents an elegant circumvention of the biophysical restrictions of transcribing such a large gene, the fact nevertheless remains that under most circumstances, demand for dp427 mRNA is indeed low: protein levels are sufficient and stable, with turnover being slow. Under normal, healthy conditions, dystrophin transcription thus appears to be continuously wasteful for no meaningful benefit. This inefficiency is demonstrably tolerable (and indeed given the energetic demands of healthy muscle, even a near-constant commitment to the 2.3 million bases of dystrophin transcription likely represents only a tiny fraction of total myofibre metabolism), but given that dystrophin is an ancient gene that predates mammals substantially [15], the continued retention of such a transcriptional system implies that it remains of situational use.

As we have suggested previously [7, 10], there are two notable scenarios where dystrophin protein (and thus dystrophin mRNA) might be in high demand: muscle development during embryogenesis and muscle repair following injury. The former establishes myotubes *de novo*, while the latter generates new myotubes to either replace lost fibres, or reconnect undamaged regions of established myofibres (segmental repair). In all these cases, the growing sarcolemma of these nascent myotubes begins in an essentially dystrophin-negative state.

## Muscle injury

Skeletal muscle tissue is highly regenerative: provided the ECM architecture is preserved, muscle can repair completely following even near-total myofibre destruction, using a dedicated population of stem cells (satellite cells, SCs) to mediate this repair. In mature muscle these cells lie dormant within the SC niche: beneath the basal lamina, but outside the sarcolemma. Quiescent SCs are challenging to distinguish histologically from adjacent myonuclei, but they can readily be recognised immunohistochemically by expression of the paired-homeobox gene *Pax7*. Following muscle damage, these cells activate, exit the SC niche and proliferate. Early proliferation events can be asymmetric: one daughter lineage returns to the niche and quiescence to maintain the stem cell population, while the other lineage remains proliferative and migrates to the site of injury [16, 17]. Clearance of damaged/destroyed tissue by scavenging macrophages occurs early and is essential for efficient subsequent regeneration, as demonstrated in elegant recent work by Collins *et al* [18], using inducible *Pax7-GFP* systems to study muscle regeneration following barium chloride (BaCl_2_) injury: within the newly vacated ECM, activated SCs undergo rapid proliferation and differentiation to give rise to large numbers of close-packed myoblasts. These myoblasts then align and fuse in a coordinated wave, either with injured but viable myofibres to restore integrity, or (where fibres have been destroyed in their entirety) to each other, to form long multinucleated myotubes. These nascent myotubes are histologically distinctive, with minimal sarcoplasm and many close-packed myonuclei aligned centrally. As the fibres hypertrophy and mature within the ECM network to restore muscle architecture, a second wave of fusion events contributes additional nuclei: these secondary nuclei are incorporated peripherally, while the initial centrally located myonuclei remain at the centre of the maturing myofibre even after repair is complete [18, 19]. In larger mammals these nuclei do ultimately migrate to the periphery, but in mice this central nucleation persists apparently indefinitely, serving as a permanent histological marker of regeneration (see Figure 1D). These regenerative stages can also be captured transcriptionally, as repair recapitulates many features of embryonic muscle development: myoblast differentiation involves the same progression of myogenic basic helix-loop-helix (bHLH) transcription factors (*Myf5*, *MyoD* and *Myogenin*), and developing myotubes similarly first express embryonic and then neonatal myosin heavy chains (*MYH3* and *MYH8,* respectively), before eventual fibre-type fate choice [20-22].

Dystrophin itself is also expressed at several stages during the repair process, though the temporal and spatial aspects of this expression remain under-characterised. Dp427m is transiently expressed within activated SCs in a spatially-restricted manner that influences early asymmetric division [17], but is not found in proliferating myoblasts. Indeed, the 16 hour transcription is likely mutually exclusive with cell division, as we have noted previously [9]: myoblasts instead express the short isoform, dp71 [23, 24]. Expression of dp427m commences only later, when myoblasts exit mitosis, fuse and differentiate to form myotubes. At this early stage, dystrophin protein would be both required but initially absent: a high demand scenario. Demand would remain high as sarcolemmal area increases throughout myotube maturation and hypertrophy, only returning to a basal (low demand) state once muscle repair is essentially complete. Under our transcriptional model, we hypothesise that cells might initiate dystrophin transcription early (potentially as soon as exiting the mitotic program, and prior to any overt requirements at the sarcolemma) - the long lag between initiation and completion cannot be circumvented, and thus early ‘presumptive’ commitment to expression minimises downstream delays. We further suggest that a system inherently optimised for essentially continuous delivery would ensure that subsequent supply of dystrophin mRNA remains uninterrupted. Placing control at the post-transcriptional level would allow this constant supply to then be titrated closely to need. Stabilizing the majority of completed transcripts would result in rapid accumulation of dp427 mRNA, greatly increasing translational output while demand for protein remains high; once demand returns to low turnover/maintenance levels, surplus mature transcripts (and most future transcripts) could be instead degraded, preventing accumulation of unwanted dystrophin at both mRNA and protein level. In essence, early activation of a strong, “once on, always on” promoter, combined with post-transcriptional fine-tuning, might be the only way to meet (but not subsequently exceed) the high demand in these rare, transient but critical scenarios. In support of this hypothesis, we note that Tennyson and colleagues reported longer transcript half-lives in differentiating myogenic cell cultures [6]; similarly, we typically observe greater numbers of mature mRNAs via smFISH in embryonic myotubes [9, 10]. Dystrophin expression in embryogenesis however involves multiple isoforms, found at different stages and in different tissues. Instead, here we test our theory empirically within a more controlled model system: investigating expression of dystrophin mRNA within injured skeletal muscle throughout the repair process.

## Methods

### Ethics statement

24 wild-type mice (strain C57BL/10J) were used for this longitudinal study, of mixed sexes (9 male, 15 female), all 5-7 months of age. Mice were bred and used under UK Home Office Project Licence PPL 70/7777, approved by the Royal Veterinary College Animal Welfare and Ethical Review Board. All mice were held in open top cages in a minimal disease unit at an average 21°C in a 12 hours light/ 12 hours dark light cycle with food and water provided ad-lib.

### Study design and sample collection

Five time points were used to follow muscle regeneration: 2, 4, 7, 14 and 30 days post-injury (DPI), corresponding approximately to: acute degeneration; clearance and activation of the repair programme; early regeneration; late regeneration, and complete repair, respectively. Mice were allocated to sample groups as shown in figure 1D (2 DPI, N=4; all other time points, N=5). Dystrophin expression should not differ between sexes (only one locus of this X-linked gene is transcriptionally active in females), but assignments were randomised within each sex to avoid uneven allocation of sexes between groups (2 DPI, 1 male, 3 females; all other time points, 2 males, 3 females).

Injury was elicited by BaCl_2_ (Sigma) injection into the tibialis anterior (TA) muscle using 20ul of 1.2% BaCl_2_ in sterile saline to promote focal muscle damage. All injections were performed under general anaesthesia using fentanyl/fluanisone (Hypnorm, Vetapharma, Leeds, UK) and midazolam (Hypnovel, Roche, Welwyn Garden City, UK) delivered intraperitoneally as published previously [25]. After recovery all mice were examined daily for adverse reaction to injury.

At the appropriate collection time (2, 4, 7, 14 or 30 DPI) mice were killed by cervical dislocation and muscles (the TA and the adjacent extensor digitorum longus (EDL), both injured and uninjured contralateral) were harvested rapidly post-mortem. Muscles were inspected visually to confirm injury/on-going repair: all injured muscles exhibited signs of injury consistent with the appropriate time point, no uninjured muscles appeared damaged. Muscles were then mounted in cryoMbed mounting medium (Bright) on histological corks and rapidly frozen under liquid nitrogen-cooled isopentane. Muscles were mounted in longitudinal orientation in a relaxed state (as described previously [7]) to allow longitudinal cryosectioning. To ensure complete coverage of the damaged region, both the TA and EDL muscles were mounted together. Uninjured contralateral muscles were collected and mounted similarly as controls. All samples were then stored at -80°C until use.

### Cryosectioning

Mounted muscle samples were removed from -80°C and transferred to a cryostat (OTF5000, Bright) to equilibrate to cutting temperature. Samples were cryosectioned (8µm thickness) in longitudinal orientation and mounted on glass slides (Superfrost PLUS): sections were collected serially to enable comparison between ISH and immunofluorescent labelling. To preserve RNA integrity for downstream FISH, sections were air dried at -20°C for 30-60min before storage in sealed slide boxes at -80°C. Additional sections were collected (30-40, from the same sectioning depth) into pre-chilled microcentrifuge tubes for subsequent RNA isolation.

### Histological staining

Haematoxylin and eosin staining was conducted as described previously [26]: slides were equilibrated to room temperature, then immersed in diluted Harris’ haematoxylin (1:1 distilled water) for 3 min. Staining was regressed via acid alcohol, followed by blueing under running tap water for 3 min. Slides were incubated in eosin yellowish (0.5% in water) for 3 min, washed quickly in distilled water and dehydrated through graded alcohols (70–100%). After equilibration to xylene (>1 hour) slides were mounted using DPX (Solmedia).

Acid phosphatase staining used the method described previously [27]: 200 ml sodium acetate staining solution (320mM, pH 4.8) was combined with 4 mg naphthol AS-B1 phosphate (1% stock in dimethylformamide) and mixed well. Separately, 3.2 ml pararosaniline-HCl solution (4% stock in 20% conc. HCl) was mixed dropwise with 3.2 ml sodium nitrite (4% stock in distilled water, prepared fresh), incubated at RT for 2 mins, then mixed with the staining solution. pH was returned to 4.8 by careful addition of 1M NaOH. RT equilibrated slides (as above) were immersed in staining solution at 37°C for 2-4 hours. Slides were washed in distilled water to terminate the reaction, then counterstained (1 min) in Mayer’s haemalum (Sigma), followed by bluing under running tap water for 3 min. Slides were dehydrated rapidly through graded alcohols (∼30 sec per stage) then allowed to dry completely before equilibration to xylene and mounting in DPX as above.

### Immunofluorescence

Immunofluorescence was conducted using slides equilibrated to room temperature (as above). Sections were ringed in hydrophobic barrier pen (ImmEdge, Vector labs) and blocked in 10% goat serum (in phosphate buffered saline supplemented with 0.05% tween-20 (PBS-T)) for 1 hr at RT. Primary and secondary antibody incubations were 1 hr at RT, with three washes of PBS-T in between each incubation. All antibodies were diluted in PBS-T. Primary antibodies and dilutions used were as follows: dystrophin (rabbit polyclonal Ab15277, Abcam, 1:200); perlecan (rat monoclonoal A7L6, Invitrogen, 1:1000); labelling of infiltrating mouse IgG used secondary antibody alone. Secondary antibodies were all AlexaFluor Plus (Thermofisher): anti-mouse 488, anti-rat 555 and anti-rabbit 647 (all at 1:800). Labelling of embryonic myosin used the Zenon labelling system (Thermofisher): antibody (mouse monoclonal MHCd, Novocastra) was preincubated with Alexafluor488-labelled FAb fragments (5 mins, RT) followed by blocking with non-specific IgG (5 min, RT): the resultant primary/fluorophore conjugate mix (final primary antibody dilution 1:40) was used alongside secondary antibody labelling, above. Following immunolabelling, nuclei were labelled with Hoechst 33342 (1:2000, 5 mins). Slides were mounted using ProLong Gold antifade (Thermofisher).

### Multiplex FISH

Fluorescence in-situ hybridisation (FISH) was conducted using the RNAscope V2 multiplex labelling system (ACDbio/BioTechne), as described previously [7, 9]. Slides were removed from -80°C and immediately immersed in chilled (4°C) 10% neutral buffered formalin for 1 hour. Slides were dehydrated through graded alcohols (50%, 70%, 100%), air dried and then baked at 37°C for 1 hour. All remaining steps were according to the manufacturer’s protocols for fresh frozen tissue. Slides were labelled with probes to the satellite cell marker Pax7 (Mm-Pax7-C3, Cat. No. 314181-C3), the cell division marker Ki67 (Mm-Mki67-C3, Cat. No. 416771-C3), and to dystrophin, using the multiplex probe set described previously [9, 10], namely Mm-Dmd (Cat. No. 452801), Mm-Dmd-O1-C2 (Cat. No. 529881-C2) and Mm-Dmd-O2-C3 (Cat. No. 561551-C3), recognising the 5’ (exons 2-10), middle (exons 45-51) and 3’ (exons 64-75) regions of the dystrophin dp427 transcript, respectively. Unless indicated otherwise, all C1 probes were labelled with TSA-Cy3, all C2 probes with TSA-Cy5, and all C3 probes with TSA-Opal 520 (all TSA dyes supplied by Akoya biosciences). Nuclear labelling used Hoechst 33342 (1:2000 dilution, 5 mins). Slides were mounted using ProLong Gold as above.

### Imaging

Brightfield images (H&E, Acid phosphatase) were collected via a DM4000B upright microscope (Leica Microsystems, Milton Keynes, UK) using either a 10x or 20x objective (HC PL FLUOTAR PH1/PH2, NA=0.3/0.5, respectively). Individual fluorescence images (ISH, IHC) were collected using a Nikon Eclipse Ni-E microscope (objective: 20x PLAN APO, NA=0.8) with D-LEDI light source (DAPI/GFP/TRITC/Cy5 filter cubes -Semrock). Whole section fluorescence images were prepared using a DMI6000 (Leica) with a motorised stage, using a EL6000 light source (A4, L5, N3 and Y5 filter cubes - Leica)-multiple images spanning the entire section were collected at 20x (HCX PL FLUOTAR PH2, NA=0.5), and then background corrected and merged using LASX software (Leica).

### Image analysis

Quantitative analysis of Dystrophin 5’ ISH probe intensity was conducted as described previously [7]: low-exposure (non-saturating) images were collected (3-4 imaging fields per section), and nuclear/sarcoplasmic foci (100-300 of each per image) were then manually defined as regions of interest (ROIs) using ImageJ. Total (background-subtracted) 5’ fluorescence intensity of each ROI was then measured. Sarcoplasmic intensity values (corresponding to individual mRNAs) were used to determine mean per-transcript fluorescence for each ISH experiment: these values were used to estimate corresponding nascent transcript numbers per nucleus. Images were collected from sections prepared from three injured (14 DPI), and three uninjured muscles. A subset of images was also collected from undamaged regions of one injured muscle.

### RNA isolation and cDNA synthesis

RNA was isolated from frozen muscle cryosections using TRIzol (Invitrogen) as described previously [28], with inclusion of an additional chloroform extraction (1:1) after the phase separation step and inclusion of 10µg glycogen during precipitation to maximise RNA yield. RNA yield and purity was assessed via Nanodrop (ND1000) and samples with 260/230 ratios below 1.7 were subjected to a second precipitation. All cDNA was prepared using the RTnanoscript2 kit (Primerdesign), using 1600ng of RNA per reaction, with oligo dT and random nonamer priming. For qPCR, 10µl of each reaction was subsequently diluted 1/20 with nuclease-free water to minimise downstream PCR inhibition. The remaining 10µl of each reaction was left undiluted to allow accurate quantification of low-abundance dystrophin mRNA via digital droplet PCR (ddPCR).

### qPCR and ddPCR

qPCR reactions were performed in duplicate (reference genes) or triplicate (all others) in 10μl volumes using PrecisionPLUS SYBR green qPCR mastermix (Primerdesign). 2µl diluted cDNA was used per reaction (∼8ng cDNA per well assuming 1:1 conversion), and PCR was conducted in a CFX384 lightcycler, with a melt curve included as standard. All Cq values were determined via linear regression. Primers to reference genes (*ACTB*, *CSNK2A2*, *AP3D1*, appropriate for healthy/damaged muscle, as determined previously [29]) were taken from the geNorm and geNorm PLUS kits (Primerdesign): all give efficiencies of 95-105% and produce single amplicons. Sequences are proprietary but anchor nucleotides and context sequences can be provided on request. Primers to *MYH3* (embryonic myosin heavy chain) and *MYH8* (neonatal myosin heavy chain) were taken from Zhou *et al* [30]. Primers to specific regions of the dp427m dystrophin transcript and to dystrophin isoform dp71 (targeting the unique first exon) were those used previously [7, 31], while primers to *SPP1* (osteopontin), *Ki67* and *Pax7* were designed to the corresponding Ensembl sequences using Primer3 (https://primer3.ut.ee/). All primers spanned large introns where possible to avoid amplification of genomic DNA. Primer sequences (where available) are provided below:

MYH3 F 5’-CTTCACCTCTAGCCGGATGGT -3’

MYH3 R 5’-AATTGTCAGGAGCCACGAAAAT-3’

MYH8 F 5’-CAGGAGCAGGAATGATGCTCTGAG -3’

MYH8 R 5’-AGTTCCTCAAACTTTCAGCAGCCAA -3’

Dp427m Ex1 F 5’-TCTCATCGTACCTAAGCCTCC-3’

Dp427m Ex3 R 5’-GAGGCGTTTTCCATCCTGC-3’

Dp427m Ex44 F 5’-TGGCTGAATGAAGTTGAACAGT-3’

Dp427m Ex45 R 5’-CCGCAGACTCAAGCTTCCTA-3’

Dp427m Ex62 F 5’-AGCCATCTCACCAAACAAAGT-3’

Dp427m Ex64 R 5’-ACGCGGAGAACCTGACATTA-3’

Dp71 F 5’-GTGAAACCCTTACAACCATGAG-3’

Dp71 R 5’-CTTCTGGAGCCTTCTGAGC-3’

SPP1 F 5’-ATCTCCTTGCGCCACAGAAT-3’

SPP1 R 5’-AGCAGTGACGGTCTCATCAG-3’

Pax7 F 5’-TGCCCTCAGTGAGTTCGATT-3’

Pax7 R 5’-GAGGTCGGGTTCTGATTCCA-3’

Ki67 F 5’-ACATTCGTATCCAGCTGCCT-3’

Ki67 R 5’-GGCTTGCTTCCATCCTCATG-3’

All data were linearised to relative quantities (RQ), and all genes of interest (GOIs) were normalised to a three-gene normalisation factor (geometric mean of the RQs of *ACTB*, *CSNK2A2*, *AP3D1*). Data were subsequently converted to log_10_(normalised RQ) for statistical analysis and graphical display.

Digital droplet PCR was conducted using 1µl of undiluted cDNA, as described above (equivalent to ∼80ng cDNA per reaction, assuming 1:1 conversion). ddPCRs were performed using Evagreen supermix (Biorad), with all primers at 100nM, using a QX200 droplet reader (Biorad). Absolute values were obtained for three regions of dystrophin dp427m and for dystrophin isoform dp71. ddPCR was conducted on a subset of samples, which were calibrated with equivalent sample qPCR Cq values to obtain standard curves for each amplicon. R^2^ values for standard curves were typically >0.99, allowing derivation of absolute values for the entire sample set (values were then normalised for variation in [cDNA] using the three-gene reference normalisation factor as above).

### Sequence analysis

Sequences for dp427m promoter regions (*Mus musculus, Homo sapiens, Canis familiaris, Equus caballus, Macaca mulatta*) were obtained from the EBI Ensembl repository (ensembl.org [32]) in FASTA format and aligned using ClustalW. Transcription factor binding sites were predicted using TFBSpred [33], using mouse and human sequence, and mapped onto the consensus alignment. Consensus TF binding motifs were extracted from the JASPAR repository [34].

### Statistical analysis

qPCR data was analysed using a Linear Mixed Model in IBM SPSS Statistics (version 29): treatment group, time point, and time/treatment (interaction) were set as fixed effects, with individual animals as random effects. Data were then subjected to post-hoc pairwise analysis, and corrected for multiple comparisons via the Holm-Šídák method (using Graphpad Prism). All data plots were prepared using Graphpad Prism. Calculations of mature/nascent transcript numbers were conducted using the methodology of Tennyson *et al* [1, 6], with the following equation:

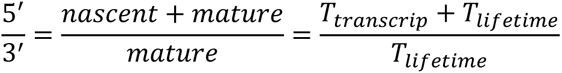

Full transcription of dp427 requires ∼16 hours from 5’ start to 3’ terminus. Transcription time used for ddPCR primer pairs was thus calibrated based on target sequence position (assuming extension rates of 40bases.sec^-1^): dp427m exon 1-3 sequence (5’) is fully transcribed ∼2 hours after initiation, while exon 62-64 sequence (3’) is completed ∼15 hours after initiation, thus T_transcrip_ = 13 hours for ddPCR. T_lifetime_ is the mean lifetime of a mature transcript: equivalent to transcript half-life divided by the natural log of 2 (i.e.T_1/2_ = T_lifetime_ *0.693).

## Results

### Muscle repair following injury

To assess repair status at each time point, we examined muscle sections histologically. Haematoxylin and Eosin staining (figure 2A) of muscles at 2 DPI was consistent with acute BaCl_2_ damage: gross myofibre architecture remained, but fibres were hypercontracted and largely devoid of myonuclei, and tissue as a whole was oedematous. Rare undamaged fibres were nevertheless present at the periphery of the injury site, and minimal inflammatory cell infiltratration was evident. At 4 DPI, cellular infiltration was prominent, with large patches of inflammation-associated mononuclear cells found throughout the tissue, scavenging and clearing damaged myofibrillar material (presumably alongside activated satellite cells and early proliferating myoblasts). By 7 DPI, widespread inflammation remained, but nascent myotubes were also visible, rendered prominent by virtue of their eosin-rich staining and multinucleate character. At 14 DPI, inflammation was markedly reduced while repair was demonstrably underway, with myotubes present in large numbers alongside more mature regenerated myofibres, identifiable by centralised nuclei. In some samples, both centrally nucleated (regenerated) fibres and peripherally nucleated (undamaged) fibres were found in close association, indicating nearly completed repair and restoration of muscle architecture. By 30 DPI, inflammation was minimal and repair was essentially complete, with all regenerated myofibres retaining long chains of central nuclei. Staining for acid phosphatase activity (a lysosome-associated inflammatory/regenerative marker [28, 35]) similarly confirmed these findings (figure 2B).

**Figure 2:**
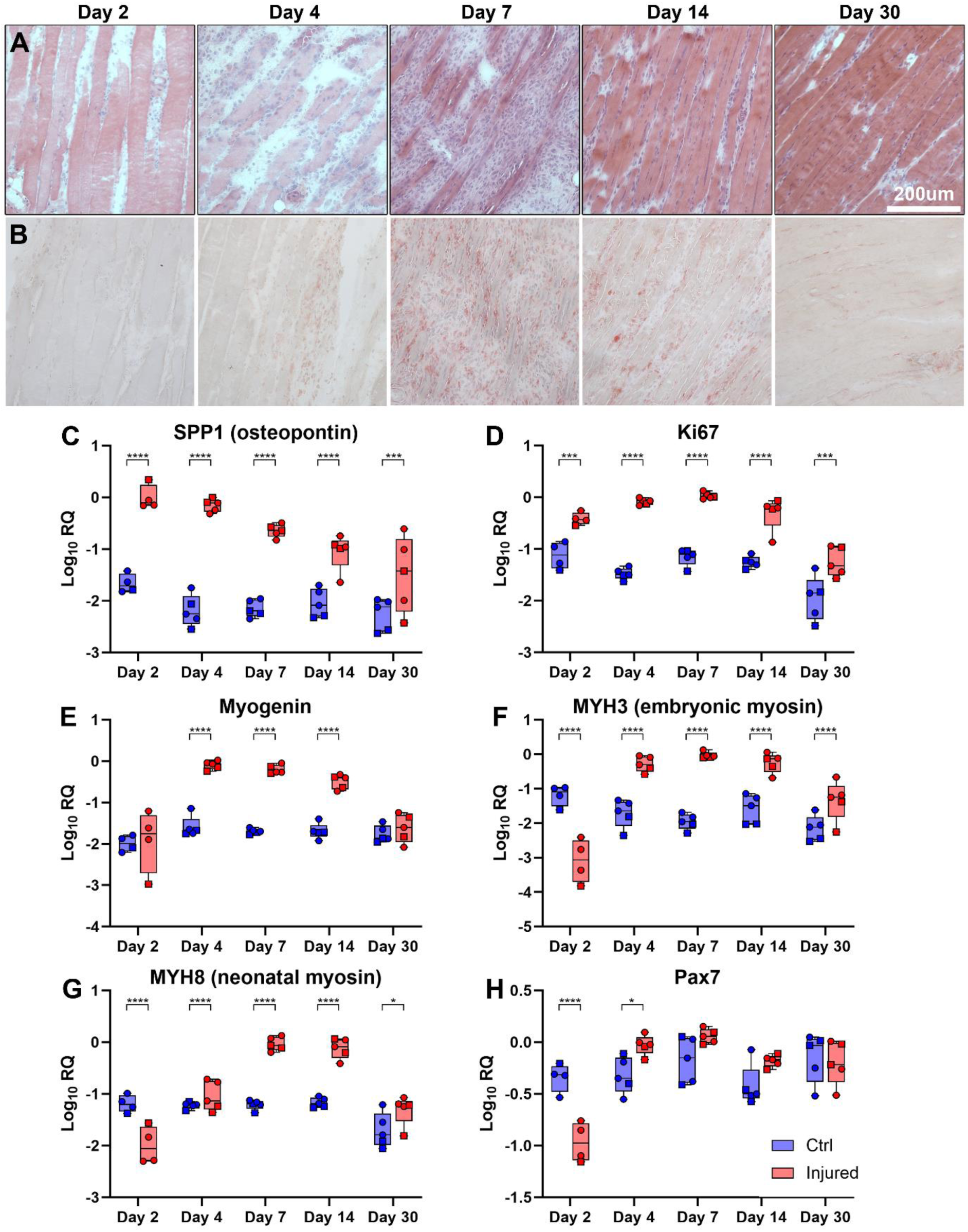
Histological and gene expression analysis of repair timeline shows key regenerative milestones. (A) Haematoxylin and Eosin (H&E) staining of injured muscle at the indicated days post injury (DPI). At 2 DPI, muscle fibres are pale, fragmented and anucleated, consistent with myonecrosis. By 4 DPI, remaining fibrillar material is being cleared by scavenging mononuclear cells, found both adjacent to, and within, dead fibres. By 7 DPI (early regeneration) nascent multinucleated myotubes are visible, surrounded by large numbers of mononuclear cells. At 14 DPI (late regeneration) myotubes have matured to centrally nucleated myofibres, which continue to hypertrophy. By 30 DPI, the repair process is essentially complete, but regenerated fibres can still be recognised by central nuclei. Staining for acid phosphatase activity (B) reveals a peak in staining from 4-14 DPI, consistent with tissue clearance and remodelling during regeneration. Scalebar: 200µm. (C-H) qPCR measurement of gene expression shows early increases in the inflammatory marker osteopontin (SPP1, C), with subsequent increases in the proliferative marker Ki67 (D) and the myogenic transcription factor myogenin (E). Expression of the regeneration-associated embryonic (MYH3, F) and neonatal (MYH8, G) myosin heavy chains is consistent with the sequential appearance of these isoforms during regeneration, while expression of the satellite cell marker Pax7 (H) increases modestly during early regeneration. All qPCR analysis used a linear mixed model, with treatment and time point as fixed variables. There was a significant effect of time (P<0.0001) for all markers, and a significant effect of treatment (P<0.0001) for all markers except Pax7 (P=0.7). Brackets on individual plots indicate post-hoc multiple comparisons testing between injured/uninjured muscles at the indicated days post injury (*=P<0.05; **=P<0.01;***=P<0.001; ****=P<0.0001).

Using qPCR, we also assessed transcriptional changes within injured muscles, measuring expression of several key genes that characterize aspects of the damage/repair process. mRNA levels of the inflammatory marker *SPP1*/osteopontin (figure 2C) were dramatically increased following BaCl_2_ treatment, commensurate with the influx of pro-inflammatory scavenging macrophages: expression at 2- and 4 DPI was almost 100-fold greater than in uninjured contralateral muscles, and though expression subsequently declined, *SPP1* remained modestly elevated even at day 30. Expression within uninjured contralateral muscles was uniformly low throughout, indicating *SPP1* expression was local, rather than systemic. mRNA for the mitotic marker *Ki67* also increased markedly following injury (figure 2D), presumably indicating proliferation of infiltrating macrophage populations. Here, however, expression further increased from day 2 to day 7 (when levels of *SPP1* were declining), potentially reflecting a shift in proliferation from scavenging macrophages to myoblasts contributing to muscle repair. Supporting this, expression of the myogenic transcription factor *myogenin* (figure 2E), and the regenerative myosin heavy chains *MYH3* and *MYH8* (embryonic and neonatal, respectively, figures 2F, G) increased dramatically from day 4 to day 14. *Myogenin* levels were highest at 4 DPI, and subsequently declined to basal levels by 30 DPI, while expression of the myosin heavy chains was highest at 7-14 DPI (with increases in *MYH3* occurring earlier than *MYH8*, consistent with the sequential appearance of these myosin heavy chains during the repair process). The satellite cell marker *Pax7* also exhibited modest increases in expression over this period (figure 2H), indicating activation and proliferation of the dedicated muscle stem cell population (expression of this marker was nevertheless low, as might be expected for a transcription factor expressed only in a minority population of cells). Interestingly, at 2 DPI, levels of *MYH3*, *MYH8* and *Pax7* were lower in injured muscles than in uninjured, possibly reflecting the stark loss of muscle tissue relative to other resident cell populations at this stage.

Taken together, these data were consistent with the histological presentations: acute degeneration and inflammation at 2-4 DPI, commensurate regeneration initiating from day 4-7, established regeneration and repair by 14 DPI, and near-complete repair by 30 DPI (though note that several gene expression markers remained modestly, but significantly, elevated even at day 30). The overall repair timeline here was fractionally slower than that reported by others [18, 21, 22] (possibly reflecting differences in animal age) but otherwise comparable.

### Dystrophin protein expression during repair: immunofluorescence

We next used immunolabelling to assess dystrophin protein during muscle regeneration, alongside other protein markers of the repair process: the basement membrane marker perlecan (for underlying tissue architecture); mouse immunoglobulin (for loss of myofibre membrane integrity); embryonic myosin heavy chain (for muscle regeneration). Perlecan revealed myofibre peripheries even during the degenerative/inflammatory phase at 2-4 DPI (figure 3A, supplementary figure 1), but sarcolemmal dystrophin protein signal within dead/damaged fibres was essentially absent. Similarly, these fibres exhibited marked influx of circulatory immunoglobulins, indicating profound membrane rupture (inset panels i). Robust sarcolemmal dystrophin staining was found in rare peripheral fibres negative for sarcoplasmic IgG (i.e. fibres spared BaCl_2_ damage, figure 3A, inset panels ii) but dystrophin protein associated with regeneration was absent at these early stages of repair. During active myotube formation at 7 DPI (with concomitant embryonic myosin expression) modest levels of newly produced sarcolemmal dystrophin were also found, though not within all myotubes (figure 3B, and inset panels iii and iv). Dystrophin deposition at the myotube membrane was moreover patchy and non-uniform at this stage, indicating as-yet incomplete restoration. By 14 DPI (figure 3C) sarcolemmal dystrophin levels associated with regenerating (centrally nucleated) myofibres appeared to be approaching healthy levels, with the strongest staining found within fibres staining only weakly for embryonic myosin (more mature fibres -inset panels v and vi). Finally at 30 DPI when the repair process was essentially complete (figure 3D), sarcolemmal dystrophin in regenerated fibres was -despite persistent central nucleation-comparable to that within adjacent undamaged fibres (inset panels vii and viii), and to that of uninjured muscle (figure 3E). Restoration of sarcolemmal dystrophin protein thus begins early in myotube development (shortly after expression of contractile proteins) and persists throughout the repair process, with levels of sarcolemmal dystrophin progressively increasing as myofibres approach maturity. Over the course of this repair, dystrophin mRNA would presumably be in near-continuous demand, and under our model this demand would be reflected in mature mRNA levels substantially above those found in healthy myofibres.

**Figure 3:**
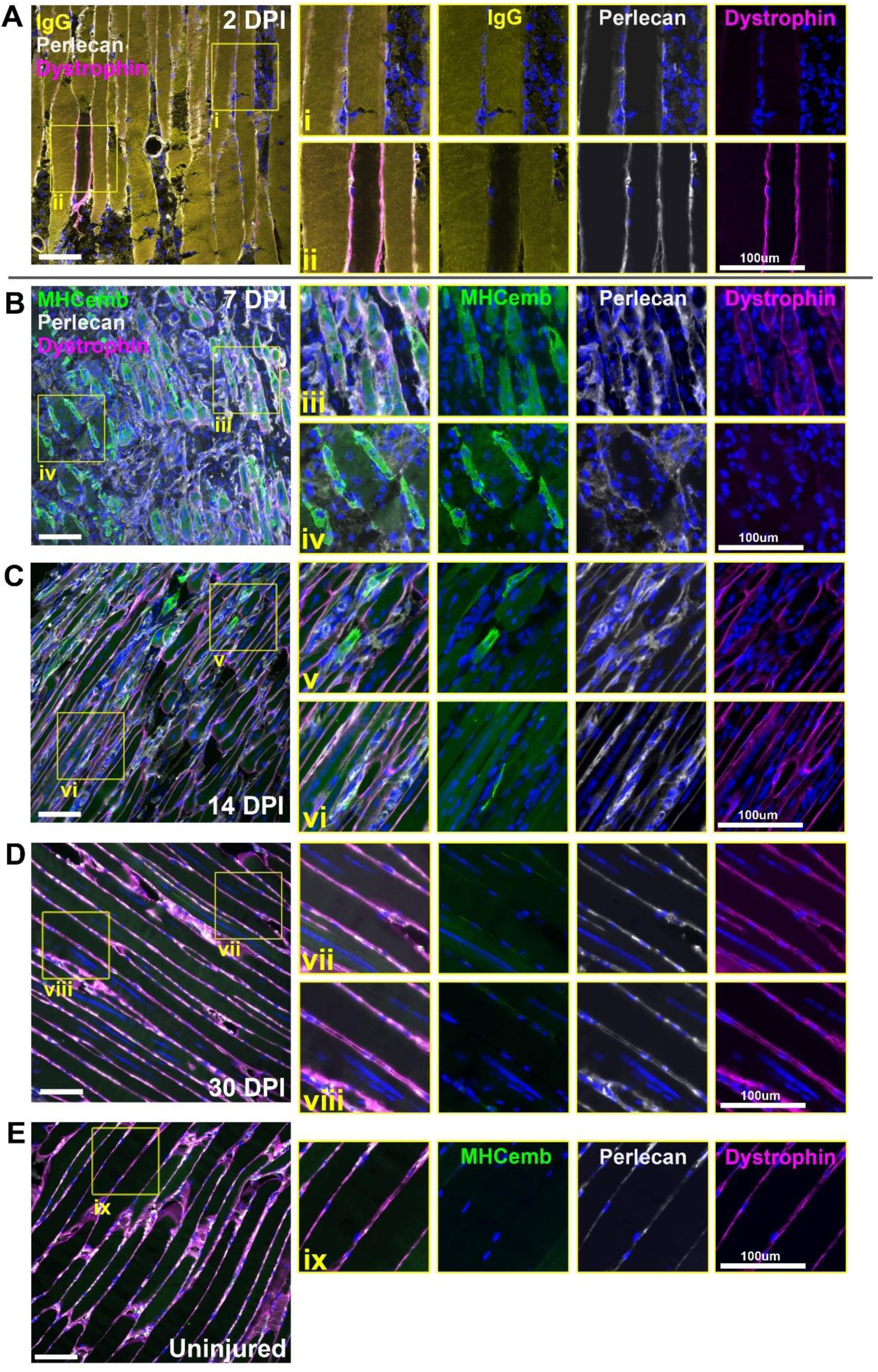
Immunofluorescence of dystrophin protein shows progressive sarcolemmal restoration throughout repair. At 2 DPI (A), sarcolemmal dystrophin staining is absent from all except rare, isolated fibres spared damage (inset ii). These fibres concomitantly are negative for infiltrating IgG, which is otherwise widespread (inset i). At 7 DPI, nascent myotubes can be identified by presence of embryonic myosin (B). Some myotubes exhibit weak/patchy staining for sarcolemmal dystrophin (inset iii) while others do not (inset iv). By 14 DPI (C) robust sarcolemmal dystrophin signal is present in many fibres. The strongest staining is found in more mature fibres that stain only weakly for embryonic myosin signal (inset vi) while fibres retaining strong embryonic myosin staining (younger fibres) show lower dystrophin signal (inset v). By 30 DPI (D), strong sarcolemmal dystrophin staining is found in all fibres, at levels comparable to uninjured muscle (E). Scalebars: 100µm

### Dystrophin mRNA expression during repair: multiplex FISH

To investigate transcriptional behaviour of dystrophin during muscle regeneration, we employed multiplex smFISH, using probes to the 5’, middle and 3’ regions of the dp427m transcript (as shown previously [7, 9]). Multiplex labelling within uninjured (contralateral) muscles (figure 4A) was consistent with our stable transcriptional model outlined in figure 1C, with a high density of nascent transcripts within myonuclei, and more modest numbers of punctate mature sarcoplasmic mRNAs, predominantly localized near the sarcolemma. Labelling within injured muscle, conversely, revealed dramatic stage-specific differences. In samples collected at 2 DPI, dystrophin labelling was effectively absent for all three probes (supplementary figure 2), consistent with complete loss of expression within dead/dying myonuclei. By 4 DPI, however, rare dystrophin-positive foci were detectable within small, typically peripheral, regions (figure 4B). These foci were exclusively nuclear, with signal concomitantly near-exclusively 5’ and middle probe only, indicating nascent, incomplete transcripts: new loci of dystrophin transcription, either within activated satellite cells undergoing asymmetric division, or within fusing myoblasts/early myotubes (absence of mature mRNA also agrees with the lack of detectable dystrophin protein at 4 DPI). At 7 DPI (when dystrophin protein was detectable within nascent myotubes) this situation was starkly altered: intense nuclear signals of 5’ and middle probe were found within the distinctive centrally located myonuclei of newly formed myotubes, indicating high numbers of nascent mRNAs (figure 4C). These myotubes were also host to markedly higher numbers of mature transcripts, with many foci of all three probes found within the sarcoplasm (substantially more than were present in uninjured fibres -compare figures 4A and C). Accumulations of nuclei labelled prominently with punctate 3’ foci only could also be found (figure 4C, inset panel iii) indicating expression of dp71, here presumably reflecting either endothelia involved in repair-associated angiogenesis, or myoblasts (which transiently express this short dystrophin isoform [23]). By 14 DPI, the central nuclei of maturing myofibres remained strongly positive for nascent transcripts, and the growing sarcoplasm was still richly populated with punctate signals from all three probes (figure 4D). This dramatic increase in mature transcript numbers was clearly repair-associated: where both regenerating and undamaged fibres could be readily discerned within the same section, transcript numbers within the latter were modest (figure 4D, inset panel viii). Finally, by 30 DPI, when dystrophin signal at the protein level suggested near-complete restoration, transcriptional differences had largely stabilised: myonuclear 5’ and middle probe foci within both peripherally nucleated (uninjured) and centrally nucleated (repaired) fibres were of similar sizes and intensities, and sarcoplasmic transcript numbers were also comparable (figure 4E).

**Figure 4:**
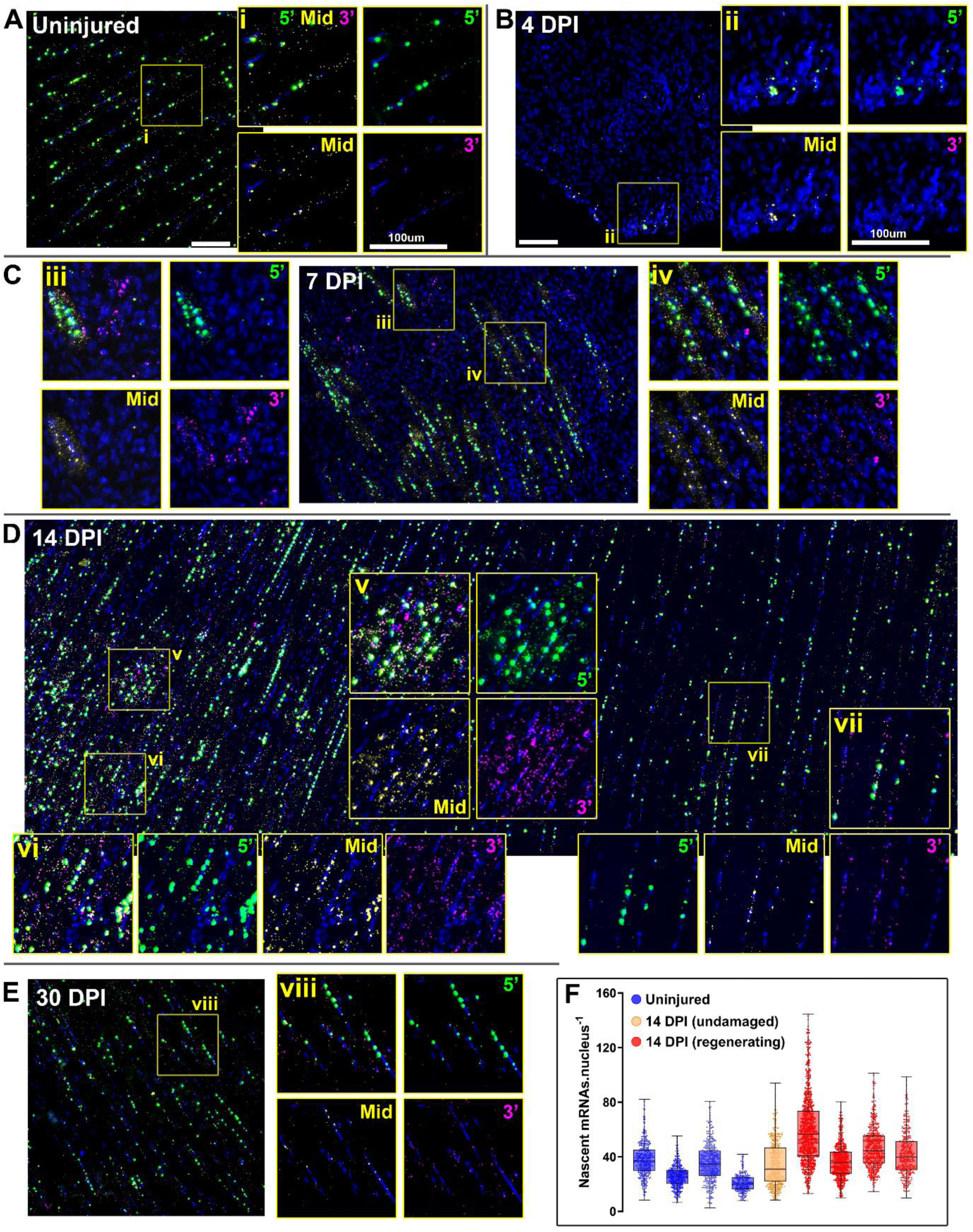
Multiplex ISH of dystrophin mRNA reveals increased transcription and mRNA stability during repair. Multiplex labelling of dystrophin mRNA in uninjured muscle produces a characteristic pattern (A): myonuclei are host to large, intense foci of 5’ probe (green), slightly smaller foci of middle probe (yellow), and punctate foci of 3’ probe (magenta), indicating multiplex labelling of nascent dp427m transcripts at different states of maturity. Punctate sarcoplasmic foci of all three probes indicate mature transcripts, found in modest numbers and typically enriched near the sarcolemma (inset i). By 4 days post injury (B) dystrophin expression resumes, restricted only to rare patches of nuclei: labelling is predominantly 5’ and middle probe nuclear foci, with minimal 3’ probe labelling (inset ii) indicating nascent transcripts only. At 7 DPI (C), myonuclei within nascent myotubes are host to intense 5’ and middle probe foci, indicating increased transcription, while myotube sarcoplasm is richly labelled with all three probes (inset iv) indicating high numbers of mature transcripts. Clusters of nuclei labelled with 3’ probe only can also be observed (inset iii) indicating expression of dp71 in either myoblasts or endothelia. Both nascent and mature transcript levels remain elevated at 14 DPI (D), with strong myonuclear foci and many mature transcripts within growing myofibres (insets v and vi). Dystrophin expression within adjacent undamaged myofibres is however comparable to healthy muscle (inset vii), as is expression at 30 DPI (E). Quantification of myonuclear 5’ fluorescence intensity (F) allows numbers of nascent transcripts per myonucleus to be determined: each datapoint represents a single nucleus (100-300 nuclei measured per muscle). Overlaid box and whisker plot shows median and upper/lower quartiles. Myonuclei within regenerating myofibres at 14 DPI are host to approximately 50% more nascent transcripts (30-60 per nucleus) than those within healthy or undamaged fibres (20-40 per nucleus). Scalebars: 100µm

Taken together, these findings largely support our hypothesis: nascent transcription commences early in the repair process (substantially ahead of protein demands, as might be expected), and subsequent mature transcripts appear to be predominantly stabilised rather than degraded, consequently accumulating in high numbers while elevated translational demand persists. However, 5’ and middle probe signal of nascent mRNAs within immature myofibre myonuclei was markedly more intense than labelling within myonuclei in uninjured fibres (particularly evident at 14 DPI -see figure 4D, inset panels v and vi), indicating greater rates of initiation during regeneration.

To assess these increases quantitatively, we measured fluorescence intensity of nuclear foci within a subset of our 14 DPI samples, which we have shown can be used to estimate per-nucleus nascent transcript numbers [7]. Individual nuclear counts spanned a considerable range (figure 4F), as we have shown previously: lower numbers presumably reflect newly initiated transcriptional loci, while higher numbers are likely to include inadvertent measurement of overlapping foci from adjacent nuclei (particularly within inured muscle where many nuclei are closely associated). Despite this, the bulk of the myonuclei in uninjured muscle were host to 20-40 immature mRNAs: values that agree with those reported previously [7]. Mean transcript numbers within myonuclei of regenerating myofibres were conversely modestly higher (30-60 per nucleus), while nuclei within undamaged regions of injured fibres appeared similar to those in uninjured muscle. A ∼50% increase in initiation is modest in transcriptional terms (where larger fold-changes are more typical), but would nevertheless result in greater mature transcript numbers, potentially explaining at least some of the observed increases. Furthermore, the sarcoplasmic volume of immature myofibres (and especially nascent myotubes) is lower than that of mature myofibres, and these regenerating fibres are moreover typically host to greater numbers of myonuclei: the high numbers of mature transcripts might thus simply be a concentration artefact, i.e. similar transcript numbers within a smaller volume.

### Dystrophin mRNA expression during repair: qPCR

To address this question, we measured dystrophin transcript numbers via qPCR/ddPCR, using primer pairs targeted to different transcript regions (5’, exons 1-3; middle, exons 44-45; 3’, exons 62-64), in a manner analogous to our multiplex FISH. 5’ sequence, present in essentially all dp427m transcripts, thus represents total dystrophin, while 3’ sequence represents predominantly mature sequence. Accordingly, the ratio of these sequences reflects the fraction of mature transcripts, and at equilibrium this is determined solely by transcription time and mature transcript half-life, not by initiation rate (see methods). A ∼50% increase in nascent mRNAs detected via ISH would generate only equivalent increases in mature mRNAs, likely insufficient to explain the numbers of mature transcripts observed. Increases in mature transcript half-life would conversely result in marked increases in mature transcript number, regardless of transcriptional initiation rates. As shown in figure 5A, total dp427m expression within uninjured muscle remained stable regardless of time point (∼5,000-10,000 transcripts.ng RNA^-1^, similar to values reported previously [7]). Levels of middle and 3’ sequence (figure 5B, C) similarly remained unchanged, though these latter sequences were present in lower amounts as expected, reflecting, at steady state, the smaller numbers of transcripts of sufficient maturity to contain middle or 3’ sequence, respectively.

**Figure 5:**
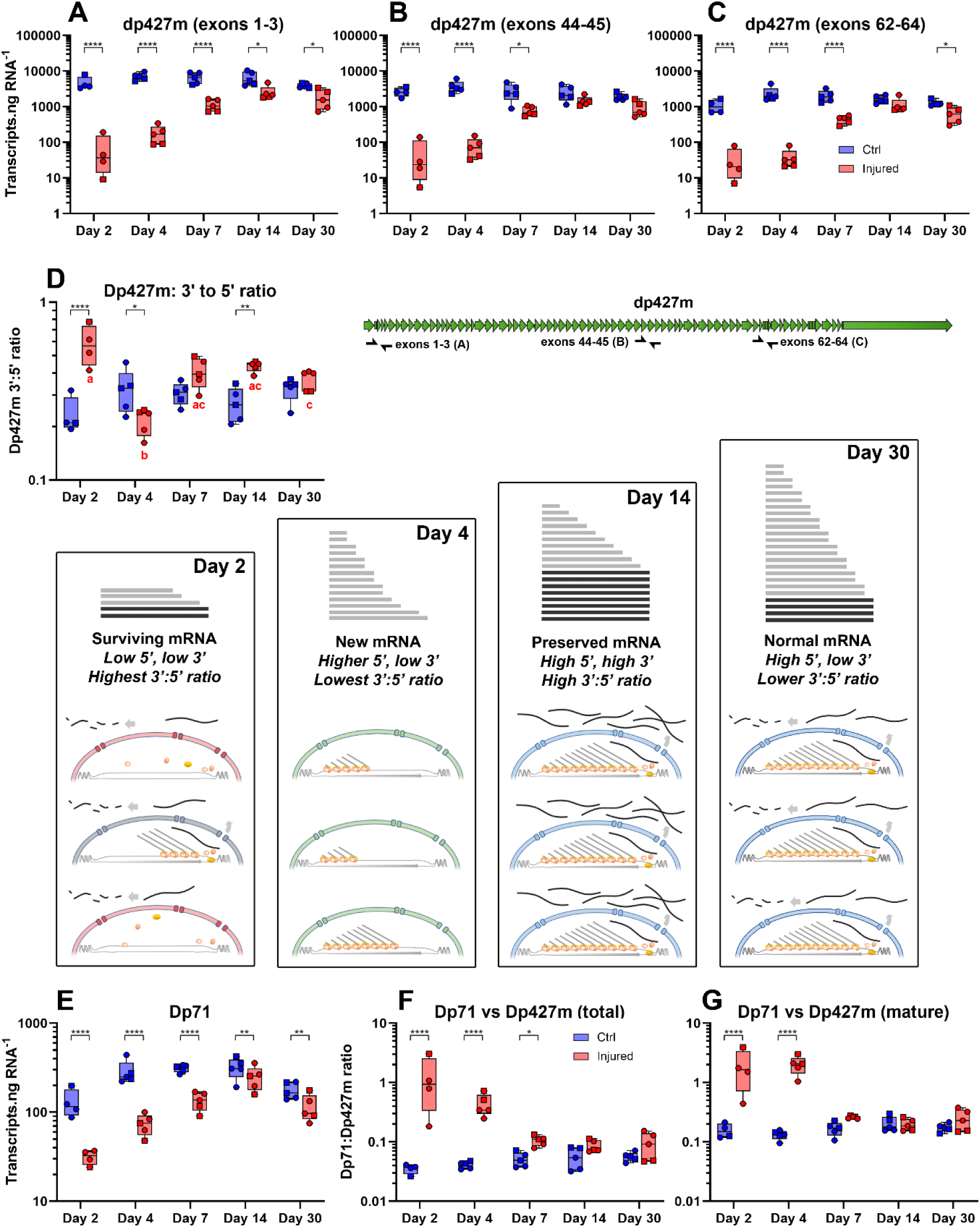
qPCR/dPCR analysis of dystrophin mRNA during repair confirms enhanced mRNA stability during repair. Absolute quantification of dystrophin mRNA in healthy and injured muscles over the course of the repair process, measuring levels of 5’ sequence (exons 1-3, A), central sequence (exons 44-45, B) and 3’ sequence (exons 62-64, C) -see transcript cartoon for approximate locations of PCR primer pairs. In uninjured muscle, transcript numbers remain comparable throughout the experiment regardless of sequence region, though individual sequence abundances (5’>>central>>3’) reflect the ratio of nascent to mature mRNAs (∼75-80% of total dp427 is nascent). In injured muscle at 2 DPI, levels of dp427 mRNA drop by 2-3 orders of magnitude regardless of sequence position, but at 4 DPI, 5’ and central sequence rise while 3’ does not. By 7 and 14 DPI, levels of all sequence regions increase, but 3’ increases more markedly than 5’ or central sequence. At 30 DPI, transcript numbers are returning to levels comparable with uninjured muscle, regardless of region. Expressing 3’ sequence (mature) as a fraction of 5’ sequence (total) highlights these transcriptional changes (D). During myonecrosis at 2 DPI, transcription ceases but rare mature dp427m transcripts remain (box ‘Day 2’) -levels of 3’ and 5’ sequence are low, but comparable. At 4 DPI, new transcripts are initiated, but are not yet complete (box ‘Day 4’), giving low 3’:5’ ratios. From 7to 14 DPI, transcription continues, and mature transcripts are stabilised (box ‘Day 14’) to supply demands, resulting in high 3’:5’ ratios. By 30 DPI demand falls and degradation of mature transcripts increases (box ‘Day 30’): 3’:5’ ratios fall once more. Expression of the short dystrophin isoform dp71 is low in skeletal muscle (E), levels decrease further following injury but recover quickly: expressed as a fraction of total dp427m (F) or mature dp427m (G), this isoform represents only 2-5% of healthy muscle dystrophin, but during early repair accounts for 50% or more of total dystrophin. Brackets on individual plots indicate post-hoc multiple comparisons testing between injured/uninjured muscles at the indicated days post injury (*=P<0.05; **=P<0.01;***=P<0.001; ****=P<0.0001). For 3’:5’ ratios, post-hoc multiple comparisons between time-points are indicated by letters, where groups sharing a letter are not significantly different: a vs b: P<0.0001; a vs c: P<0.05; no uninjured groups were significantly different at any time point).

Expression within injured muscle conversely altered substantially over the repair process: at 2 DPI, total numbers of dp427m transcripts (5’ sequence) fell dramatically (only ∼10-100 transcripts.ng RNA^-1^), matching the absence of detectable dystrophin ISH signal at this stage. Levels at 4 DPI were slightly higher (∼100-500 transcripts.ng) and continued to rise steadily over the course of the repair process, though did not return to uninjured levels even by 30 DPI. Levels of middle and 3’ sequences mirrored these changes broadly, but not exactly, remaining lower at 4 DPI, but also subsequently rising more abruptly: crucial differences that indicated changes in transcriptional behaviour.

To illustrate this more clearly, we normalized 3’ sequence (mature) to 5’ (total), a ratio that corrects for changes in overall dystrophin expression (figure 5D). Within uninjured muscle, 3’:5’ ratios varied modestly between individuals, but were typically 0.2-0.3 regardless of time point, indicating that at healthy steady state ∼20-25% of dystrophin transcripts are mature, with a mean lifetime of ∼4 hours (values that agree well with those reported previously [6, 7]). Within injured muscles at 2 DPI the ratio increased markedly over healthy levels (in some samples nearing parity) while at 4 DPI the reverse was true, with a 3’:5’ ratio significantly lower than in healthy muscle. From 7 DPI the ratio appeared to reverse again, and by 14 DPI levels of 3’ sequence relative to 5’ were significantly higher (and remarkably consistent) in all injured muscle samples, corresponding to a mean mature transcript lifetime of ∼7-8 hours. This alone would result in a doubling of mature transcript numbers independent of altered initiation rates. By day 30, ratios within injured muscles largely returned to values comparable with those of healthy muscle.

Combining these observations with absolute transcript numbers and ISH data builds a model of transcriptional dynamics during muscle injury and repair (figure 5D panels, Day 2 – Day 30). Following injury and subsequent myofibre degeneration, all myonuclear transcription (including dp427m) effectively ceases, and levels of nascent mRNAs drop precipitously: for a brief period, the only remaining dp427m mRNAs would be rare, surviving, mature transcripts. At 2 DPI, overall dystrophin transcript numbers would thus fall markedly, but the ratio of 3’:5’ sequence would paradoxically increase, exactly as observed. At 4 DPI following clearance of damaged tissue and initiation of regeneration, dystrophin transcription would begin anew, but here expression would be against a ‘blank slate’ background, with little or no mature dp427m mRNA present. At this stage, overall dystrophin mRNA levels would begin to rise, but be detected principally as 5’ sequence, not 3’, representing newly initiated transcripts that have not yet reached completion: 3’:5’ ratios would consequently drop below healthy levels. As the repair process proceeds and transcripts reach maturity (7-14 DPI), the situation would become more complex: in the dystrophin-negative environment of a newly regenerated myotube/myofibre the demand for dp427m mRNA would be high, and transcripts would thus be protected from degradation. Here dystrophin expression would increase while also shifting to a 3’:5’ ratio markedly higher than that seen in healthy muscle, reflecting the increased lifespan of protected mature dp427m transcripts (note that apparent increases in 3’:5’ ratio could also be transiently achieved by reduction in initiation, but as our ISH demonstrates, this is not the case). As myofibres reach maturity at 30 DPI and sarcolemmal dystrophin protein reaches healthy levels, the situation would normalise once more, returning to a 3’:5’ ratio consistent with excess production and post-transcriptional degradation.

Given the small but distinct patches of 3’ probe signals observed at 7 and 14 DPI (indicative of dp71 expression), we also measured expression of this short isoform. As shown in figure 5E, following injury expression of dp71 fell (and subsequently recovered) in a manner analogous to that of dp427m. As with 3’ and 5’ sequences, measurement of absolute transcript numbers further allowed these two isoforms to be directly compared across the repair process, with dp71 expressed either as a fraction of total dp427m, or as a fraction of mature dp427m. In uninjured muscle, dp71 transcripts represented only 2-5% of total dp427 (or ∼10% of mature dp427), regardless of time point. Within injured muscle, reductions in dp71 expression were less dramatic than those of dp427m, with the relative proportions of these isoforms changing throughout repair as a consequence: immediately following injury (2 and 4 DPI), numbers of dp71 and dp427m transcripts were essentially equivalent, with similar (but low) numbers of both present. Subsequent increases in dp71 expression were swifter (reflecting the short transcription time of this isoform) but also less dramatic than those of dp427m: the fraction represented by this short isoform thus remained elevated at 4 DPI, but fell over the course of regeneration. Even by day 30, however, expression appeared modestly elevated over uninjured levels (∼5-10% of total, ∼20-25% of mature), suggesting that the repair process was not fully complete even at this stage (also indicated by some other markers of damage/regeneration, above).

### Dystrophin and markers of regeneration: Pax7 and Ki67

Finally, to place our work in context within the broader regenerative landscape, we explored spatial expression of dp427 alongside more specific cell markers, combining our 5’ and 3’ dystrophin FISH probes with probes to either the SC marker *Pax7*, or the proliferation marker *Ki67*.

Dp427 was expected to partly co-localise with *Pax7* during regeneration, as asymmetric division of SCs first requires transient expression of dp427 [17]. In healthy muscle, *Pax7* labelling was as expected (figure 6A), with comparatively high signal found only within distinct, rare, peripheral nuclei consistent with quiescent satellite cells. Some signals appeared to co-localise with dystrophin labelling (both 5’ and 3’), implying a degree of co-expression even during quiescence (figure 6, inset i), however the close-packed nature of peripheral nuclei rendered this difficult to confirm (SCs are challenging to distinguish from adjacent myonuclei). Immediately following injury (2 DPI), we were unable to detect any unambiguously *Pax7* positive nuclei in any samples (supplementary figure 3): in agreement with the reduction in mRNA levels determined via qPCR (figure 2H) but perhaps unexpected from a regenerative perspective. By 4 DPI, when qPCR indicated a marked increase in *Pax7* expression, distinct patches of *Pax7*-positive cells were found closely associated with sites of nascent dystrophin expression (figure 6B), with some nuclei co-labelling with both *Pax7* and dystrophin 5’ probes (figure 6, inset iii), confirming co-expression of these genes. Similar cells were found associated with nascent myotubes at 7 DPI (figure 6C). At 14 DPI, the reestablishment of the myofibre architecture rendered *Pax7*-positive cells particularly prominent (figure 6D): these were predominantly peripheral, labelling for *Pax7* alone, or expressing both *Pax7* and nascent dystrophin, consistent with the ‘second wave’ of fusion events reported by Collins *et al* [18]. Interestingly we also detected modest punctate signals associated with (strongly dystrophin-positive) centrally located nuclei within maturing myofibres, suggesting that *Pax7* expression continues post-fusion to some extent (figure 6, inset vi). By 30 DPI, *Pax7* was once again restricted to rare, peripheral nuclei, consistent with completion of repair (as with uninjured muscle, a degree of co-localisation with dystrophin 5’ and 3’ probes was observed).

**Figure 6:**
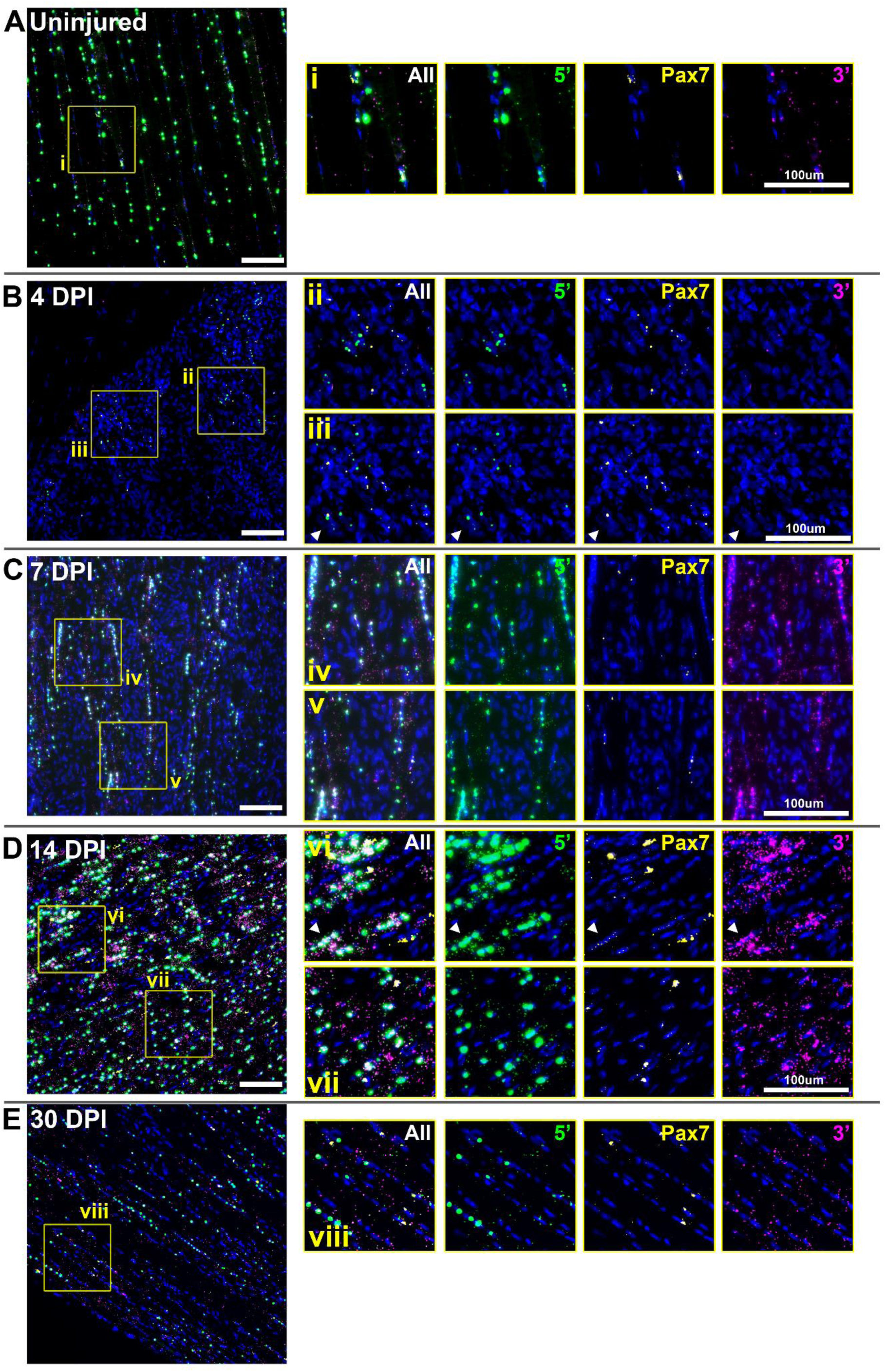
Multiplex ISH of dystrophin and *pax7* mRNA indicates overlapping expression programs. Muscle labelled with probes for dystrophin 5’ sequence (green) and 3’ sequence (magenta), alongside probes for the satellite cell marker Pax7 (yellow). In healthy muscle (A), pax7 expression is found only in rare peripheral nuclei as expected, apparently accompanied by dystrophin expression in some instances (inset i). In injured muscle at 4 DPI (B), pax7 expressing cells are typically found in close apposition to sites of nascent dystrophin expression (inset ii). Cells co-expressing dp427 and pax7 are rare but detectable (inset iii, arrowheads). Both markers remain in close proximity at 7 DPI (C, insets iv, v) though total pax7 signal appears modest. At 14 DPI (D) intensely pax7-positive nuclei can be found alongside growing myofibres, both with and without co-expression of dp427 (inset vii). Modest pax7 expression is also detectable within some robustly dp427-positive myonuclei (inset vi, arrowheads). By 30 DPI (E), pax7 expressing cells are again found residing peripheral to regenerated myofibres (inset viii). Scalebars: 100µm

Healthy muscle signal was devoid of *Ki67* expression (supplementary figure 4), consistent both with the low measured expression via qPCR (figure 2D), and the minimally mitotic nature of mature skeletal muscle. We have previously argued that dp427 might indeed be largely restricted to such post-mitotic populations purely because a 16-hour transcription time should be incompatible with active proliferation (DNA cannot be used as template for both processes simultaneously). Accordingly, we would not expect *Ki67* to co-express with dystrophin.

qPCR suggested marked increases in *Ki67* expression following injury, however ISH revealed that even within damaged, actively regenerating tissue, expression of *Ki67* was not widespread. At 2 DPI (when dystrophin expression was absent) *Ki67* positive signal was typically focal, restricted to isolated patches of mononuclear cells (figure 7A). At 4 DPI this focal distribution persisted (figure 7B): here, as with *Pax7* (above), *Ki67*-positive cells were often found in close association with sites of nascent dystrophin expression, most likely indicating proliferating SCs or myoblasts, but no unambiguously double (*Ki67*/dp427) positive cells were found at this stage. *Ki67* positive cells were similarly detected at 7 DPI, typically close to some, but not all, nascent myotubes (figure 7C, insets iv, v). Despite qPCR data indicating peak *Ki67* expression at this stage, overall ISH signal remained sparse. By 14 DPI, when muscle architecture was reestablishing, *Ki67* expression was restricted to distinct, individual nuclei located along the periphery of maturing myofibres (figure 7D). Some of these nuclei were also associated with 3’ probe, but in one instance we detected clear co-expression of *Ki67* and nascent (5’ probe) dystrophin expression (figure 7, inset vi, arrowheads). These data suggest that these peripheral nuclei were likely to be both proliferating myoblasts and SCs, and that unexpectedly, proliferation and dystrophin expression might partially overlap. Finally, by 30 DPI (figure 7E), *Ki67* expression was near-comparable to that of healthy muscle, with signal restricted to very rare individual nuclei (2-3 per entire muscle section) neither within, nor peripheral to myofibres, but associated with nuclear-rich clusters which were also often 3’ probe positive (i.e. possibly vascular endothelia).

**Figure 7:**
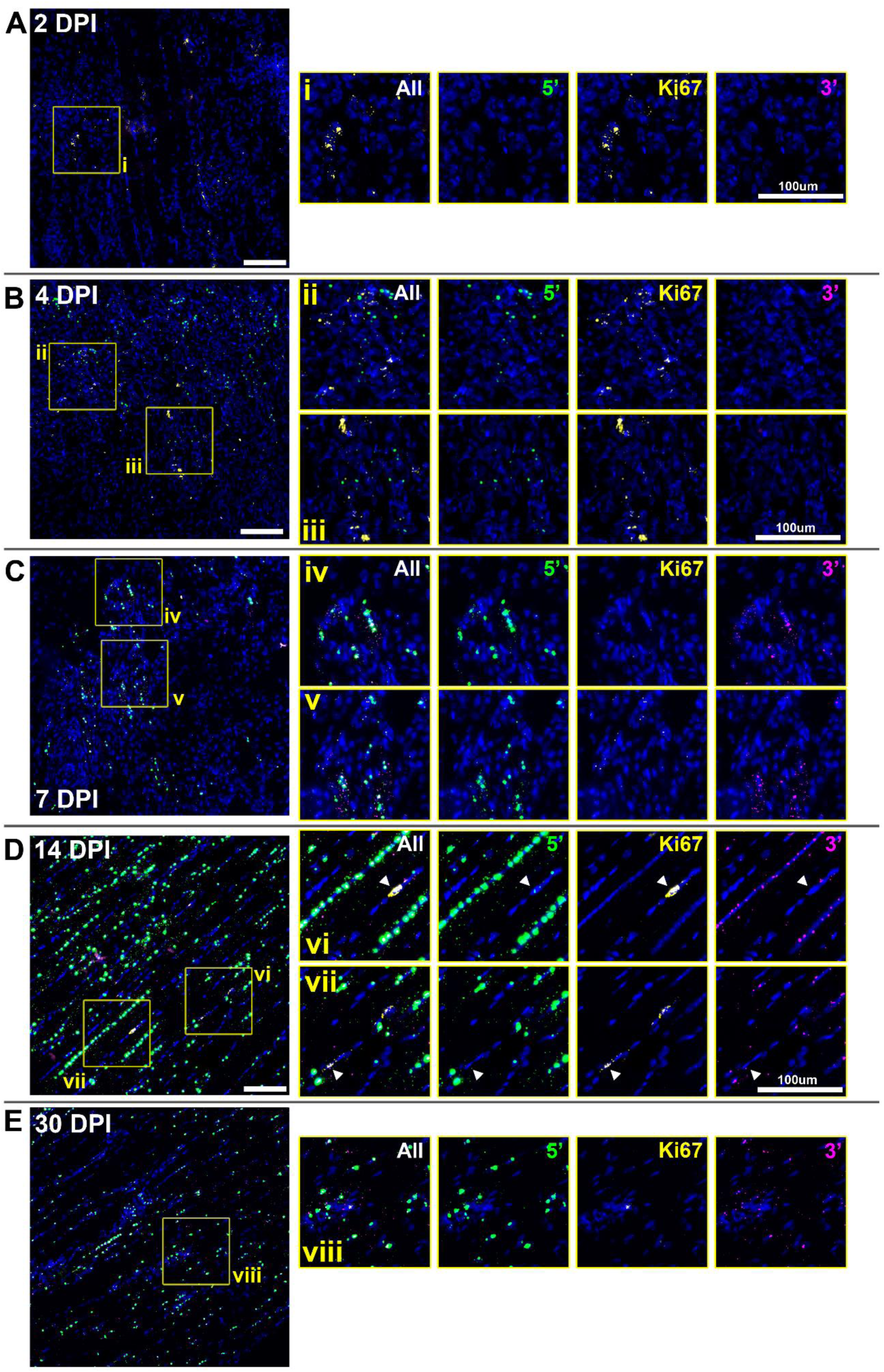
Multiplex ISH of dystrophin and ki67 mRNA reveals rare co-expression. Muscle labelled with probes for dystrophin 5’ sequence (green) and 3’ sequence (magenta), alongside probes for the cell division marker ki67 (yellow). Ki67 was detectable even at 2 DPI (A) when dp427 expression was absent, but expression was restricted to focal clusters of cells (inset i) rather than widespread. Focal ki67 clusters were also evident at 4 DPI (B), found close to patches of nascent dp427 expression, but no co-expression was detected (insets ii, iii). At 7 DPI (C), ki67 expression was restricted to rare cells found in close proximity to some, but not all developing myotubes (insets iv, v). By 14 DPI (D), ki67 was found only in peripheral (rather than central) nuclei (inset vii, arrowheads), which were negative for dp427 in all except one instance (inset vi, arrowheads) where prominent expression was found alongside punctate 5’ probe signal indicative of recently initiated dp427 transcription. At 30 DPI (E), rare Ki67 positive nuclei could be detected at fibre boundaries, in some instances co-localising with or expressed adjacent to 3’ probe signal indicating dp71 (inset viii). Scalebars: 100µm

Collectively, these data support and refine the regenerative timeline, indicating that early (day 2) proliferation is associated with non-myogenic cells, while subsequent sites of proliferation are chiefly associated with establishment of nascent myotubes and hypertrophy of maturing myofibres.

## Discussion

Expression of full-length dystrophin (dp427) is by necessity unusual: the 16-hour transcription time enforced by the enormous size of the *Dmd* gene precludes regulation via transcriptional initiation over any smaller timescale. As we have previously shown, myonuclei within multinucleate myofibres instead initiate dystrophin transcripts continuously and control mRNA levels post-transcriptionally, a wasteful but responsive ‘pay in advance’ model, ostensibly able to circumvent the otherwise unavoidable delay and meet changes in demand rapidly. In healthy muscle demand is, however, low: most mature transcripts are degraded shortly after completion, and immature transcripts represent 70-80% of total dystrophin mRNA, indicating marked oversupply. Fine control over expression might thus not only be wasteful, but unnecessary. Dp427 is also associated almost exclusively with terminally differentiated, post-mitotic cell types (principally skeletal, smooth and cardiac muscle, and neuronal lineages within the brain), lineages with differentiation processes that require multiple days, and with longer-term dystrophin demands that are both relatively low and stable. Even with a 16-hour delay, dystrophin mRNA could presumably be continuously supplied at levels sufficient to meet these modest demands. The oversupply of dp427 transcripts is nevertheless a conserved phenomenon, found in both muscle and neuronal lineages, and across multiple species (mice, dogs and humans), strongly suggesting that responsive expression remains essential (i.e. changes in demand can occur). Here we investigated dystrophin expression at both the mRNA and protein level during BaCl_2_-induced skeletal muscle injury and repair, covering key stages in the repair process over which demand for dystrophin was expected to change substantially: our findings confirm our transcriptional model but also reveal further insights into transcriptional control.

The near-complete loss of dystrophin mRNA at 2 DPI was expected, given the destruction of most myofibres. Similarly, while dead/damaged myofibres were still readily recognizable as such via basement membrane perlecan staining, dystrophin protein was absent, indicating loss/degradation of the sarcolemma. This absence of detectable protein persisted at 4 DPI, when rare patches of nuclear dystrophin 5’ probe signal (and increases in 5’ sequence via qPCR) were detected, allowing us to attribute these foci unambiguously to new transcriptional initiation. Macrophage-mediated clearance of cellular debris is essential for myotube formation [18], and at 4 DPI this scavenging was ongoing, explaining the absence of recognisable myotubes at this stage: this nascent dp427 mRNA instead appeared to be associated with mononuclear cells, and was predominantly (but not exclusively) distinct from cellular *Pax7* expression, indicating the bulk of early expression occurred within myoblasts/myocytes rather than asymmetrically dividing SCs. Use of a *Ki67* probe allowed us to assess proliferation within these cell populations, something we have previously argued might be mutually exclusive with dp427 (but not dp71) expression [9, 10], given the conflicts between DNA replication and a 16-hour transcription process. Our data here suggest this hypothesis merits revision: in at least one instance (figure 7D) the proliferation marker was unambiguously and prominently found alongside dystrophin transcriptional loci, demonstrating overlap can occur. Dystrophin expression here was however demonstrably recently initiated (small nascent foci of 5’ probe only), and *Ki67* expression chiefly increases during G_2_/M, rather than S-phase [36]: dp427 expression might thus only be precluded during S-phase, accelerating transition from proliferative to dystrophin-expressing status. Alternatively, expression might be independent of cell cycle status, but simply prematurely terminated by DNA replication, limiting successful dystrophin expression only to periods outside S-phase (given the inherently wasteful nature of basal dystrophin expression, such inefficiency is not implausible). Collectively, these data suggest that nascent dp427 mRNA expression begins remarkably early in the myogenic differentiation process, potentially coincident with exit from the cell cycle, and substantially prior to myoblast fusion. As the delay between initiation and completion must be endured at least once, committing to expression in advance of fusion would render nascent myotubes essentially ‘primed’ for completion, with the price largely paid prior to overt demands at the sarcolemma.

The following regeneration stages (7 and 14 DPI) best demonstrated these sarcolemmal demands, from formation of nascent myotubes through to maturing myofibres. Here sarcolemmal dystrophin protein was detectable, but not yet fully restored, with the continuous hypertrophy of these maturing fibres (and concomitant expansion of sarcolemmal surface area) serving to keep demand elevated. As we predicted, this increased demand resulted in marked increases in mature transcript numbers, which absolute analysis of 3’:5’ ratios confirmed was indeed achieved via increased mRNA stability. Numbers of nascent mRNAs (as represented by intensity of nuclear 5’ labelling) also increased, however, demonstrating that demand is controlled at multiple levels: over short timescales via stability, and longer periods by supply. In line with this, at 30 DPI when repair was near-complete, both stability and supply returned to near-normal levels.

We were also able to assess the dp427 expression within centrally located myonuclei, which necessarily supply all initial dystrophin demands within myotubes and immature myofibres until secondary fusion events contribute peripheral nuclei. The persistence of central nuclei in mouse muscle is unusual, but also without apparent deleterious consequence. It is not known whether these central nuclei continue to functionally contribute, however, as their location (surrounded by dense myofibrillar apparatus) might impair appropriate targeting of exported mRNAs. Others have nevertheless shown that central nuclei continue to be robustly transcriptionally active [37], and we have demonstrated nascent dystrophin expression within central nuclei of *mdx* mouse fibres [7], though the prompt post-export degradation of completed *mdx* dystrophin transcripts via nonsense-mediated decay (NMD) precluded assessment of mature mRNA behaviour. As we show here, in injured healthy muscle, high numbers of both nascent and mature transcripts were associated with central nuclei at both 7 and 14 DPI (figure 4C, D), confirming that these nuclei contribute the bulk of dystrophin mRNA to the growing sarcolemma. This expression persisted at 30 DPI, with scattered mature transcripts found both close to central nuclei and distributed throughout the sarcoplasm (figure 4E), suggesting that even after addition of multiple peripheral nuclei (which similarly express dp427) and repair completion, central nuclei continue to contribute dystrophin transcripts to the myofibre.

We acknowledge that the changes in dystrophin demand explored here represent an extreme scenario: myofibre destruction and concomitant compensatory regeneration can be induced and followed experimentally, but are unlikely to be commonplace occurrences within skeletal muscle under normal conditions (though segmental necrosis can occur following exercise [38, 39]). Milder insults however, such as membrane microtears and disruptions, likely occur on a more regular basis in healthy skeletal muscle: these might not require SC activation, but their repair would nevertheless elicit changes in demand. Membrane repair is multiphasic [40]: a ‘patch’ forms within seconds (sealing the membrane and preventing toxic calcium flux), but subsequent ‘full’ repair of the membrane architecture occurs more slowly, with the precise pathway further varying with extent of damage (including myonuclear migration [41]). Restoration of sarcolemmal dystrophin occurs during this second phase: under the model proposed here, local dp427m transcripts would be selectively preserved, allowing this restoration to begin promptly. Fine-tuning of dystrophin mRNA might be useful more broadly: hypertrophic postnatal muscle growth can exhibit circadian behaviour, and even a 24-hour cycle is challenging to match with a 16-hour transcription time: constant production coupled to post-transcriptional regulation would instead present fewer challenges.

Questions undoubtedly remain: our previous work has explored the ‘what’, demonstrating that dp427 ‘transcript imbalance’ results simply from continuous production, lengthy transcription and subsequent rapid degradation, such that most transcripts are nascent rather than mature. Our study here provides a framework for the ‘why’, showing that this counterintuitive system allows for rapid and fine control over mature mRNA levels across states of varying demand during muscle repair. The ‘how’, at present, remains under characterised. Both initiation rate and mRNA stability influence transcript levels, and while we propose stability is the principal mediator of short-term tuning, clearly both contribute. How do myofibres sense sarcolemmal dystrophin, and how is this sensing coupled to control of mRNA production and turnover? Is such nuanced behaviour regional, or generalised within a myofibre?

Increases in dp427 transcriptional initiation during early regeneration suggest involvement of stage-specific transcription factors. The dp427m promoter has multiple highly conserved binding motifs for both broad utility (*Fos*/*Jun*, *STAT*) and more muscle-specific (*SRF*, *Six4*, *TEAD*) transcription factors (figure 8A): *Six4* in particular might enhance regenerative expression of dp427, as this factor is a key activator of other early myogenic genes (including *myogenin*, which peaks early in repair -Figure 2E) [42]. Increases in expression measured here were however modest (∼50% over basal, or ∼40-60 nascent transcripts per nucleus), implying that even under peak demand, output cannot be markedly enhanced. It seems unlikely that ∼60 active transcription complexes represent even close to maximum occupancy of a gene 2.3 million bases long, thus initiation might in fact be a limiting step (also suggested by other investigators [4]). Whether this limitation takes the form of a constitutive but comparatively slow assembly of initiation complexes, or infrequent but high efficiency transcriptional ‘bursts’, remains an open question (figure 8B, C): burst expression occurs in many genes [43, 44], but the labelling and imaging approaches used here currently lack the resolution to distinguish these two possibilities.

**Figure 8:**
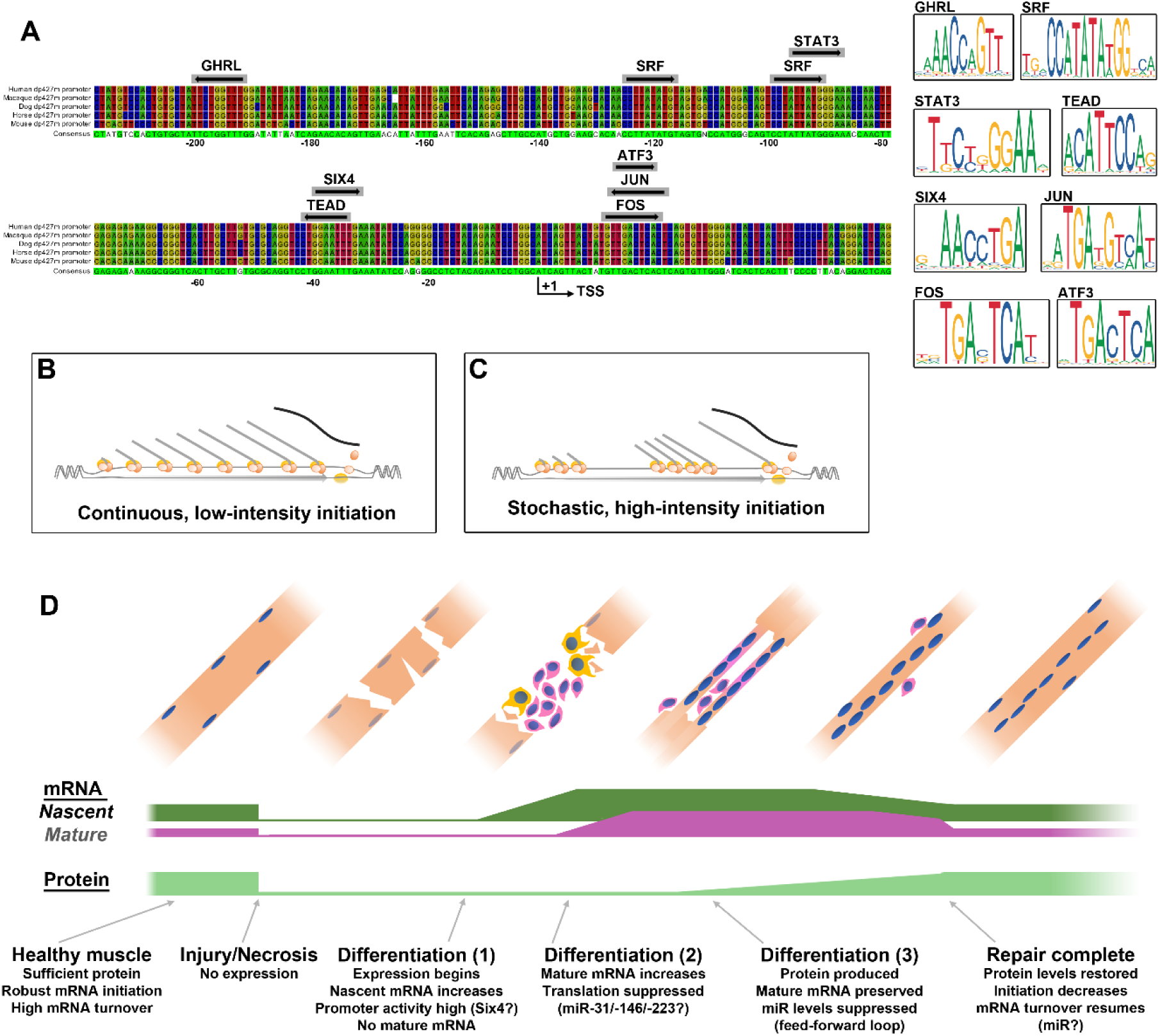
Regulation and dynamics of dystrophin expression. The proximal dp427m promoter region is highly conserved (A): alignment of human, macaque, dog, horse and mouse sequence (from ∼250bp upstream to 50bp downstream of the transcriptional start site, TSS) reveals several transcription factor binding sites of complete sequence identity. Consensus sequence is shown: green indicates universally conserved nucleotides. Predicted TF binding sites are shown, with arrow direction indicating orientation of binding motif. Consensus motif sequences (JASPAR) are shown to the right. Possible transcription modes for dystrophin expression: constitutive (B), where transcripts are initiated continuously at low levels, distributed along the <u>Dmd</u> locus; ‘bursty’ (C), where high numbers of initiation events occur at stochastic intervals, leading to ‘transcriptional bubbles’ along the <u>Dmd</u> locus, containing many transcripts of similar maturity. Timeline for dystrophin during muscle repair (D): in healthy muscle transcript degradation is rapid, and nascent mRNAs outnumber mature, while protein levels remain high and stable. During myonecrosis levels of nascent and mature mRNA, and protein all drop to near-zero. Early in differentiation, transcription begins, generating nascent mRNA numbers that surpass normal expression. As differentiation proceeds, transcripts reach maturity and are selectively preserved rather than degraded. Translation is initially suppressed through microRNA interactions, but low levels of dystrophin protein then suppress miR expression and potentiate further translation in a feed forward loop that generates sufficient dystrophin protein to supply the sarcolemma during hypertrophy. Once repair is complete, mRNA degradation resumes, and most dp427 mRNA is again nascent, not mature.

The long transcription time necessitates control via mRNA turnover for faster regulation. Our studies here confirm this occurs, implying underlying control mechanisms acting over comparable timescales. Stability, targeting and translational competence of mRNA is largely mediated through the 3’ UTR [45], and at ∼2.6kb, the dystrophin 3’ UTR is substantial. Long 3’ UTRs are inherently vulnerable to degradation [46], and (in agreement with the model here) the dystrophin 3’ UTR destabilises luciferase mRNA in myoblasts, but stabilises it in maturing myotubes [47], properties that require the well-conserved ‘Lemaire A and D’ regions at the 3’ UTR termini [48]. AU-rich elements (AREs - one or more AUUUA motifs within U-rich regions) mediate turnover in many mRNAs, directed by phosphorylation-dependent ARE-binding proteins: a responsive system. The dystrophin 3’ UTR contains only 4 conserved AUUUA sequences, however (2-3 would be expected through chance alone) and only one loosely conserved AU-rich region. A host of other proteins can interact with 3’ UTRs, influencing stability, targeting, and translation competence, however these interactions are often mediated through long and poorly conserved motifs (with secondary structure elements such as stem loops), making them difficult to identity *a priori*. The dystrophin 3’ UTR interacts with the RNA-binding protein vigilin [49], however this protein is notoriously multifunctional [50], making its role here unclear.

Many muscle mRNAs (especially long transcripts) appear to be specifically targeted to appropriate locations in a microtubule-dependent fashion [51]: given the preferential localization of dp427m transcripts near the sarcolemma, and the interactions of dp427 protein with tubulin [52], it is likely dystrophin is no exception (the excursion of transcripts exported from central nuclei supports this). This targeting does not appear to vary with demand, however, arguing against a role in turnover. Translation itself influences stability: translation and degradation factors compete for access to polyA tails [53], thus suppression of translation promotes degradation. MicroRNAs could thus play a key role: miRs can impair translation and/or promote degradation via interaction with the 3’ UTRs of target mRNAs, and can be produced swiftly, in large numbers. Multiple miRs vary during myogenesis [54, 55], promoting proliferation (miR-133) or differentiation (miR-1, miR-206) and several (including miR-31, - 146 and -223) have been shown to directly bind the dystrophin 3’ UTR and inhibit translation (blocking or oblation of these miRs also enhances protein levels in dystrophin-deficient scenarios) [56, 57]. These miRs are not reported to concomitantly enhance mRNA degradation, however: as noted previously [10] this could be partly artefactual, attributable to measurement of 5’ exon sequence for quantification (where prominent contributions from nascent transcripts would mask more subtle changes in mature mRNA numbers), but more pertinently, these miRs are also upregulated in dystrophic muscle: a dystrophin-deficient scenario where inhibition of translation should not be favoured.

These specific miRs are more likely components of a developmental feed-forward switch (as we and others have discussed [58, 59]), which would match early commitment to dystrophin expression: at initial myogenic stages, these miRs would permit dystrophin transcription while preventing aberrant, deleterious translation (dp427 protein is detrimental to cell division [60]), while post-fusion, signalling from sarcolemmal dystrophin protein would suppress miR levels and consequently potentiate translational throughput. Within a dystrophic context, this latter signalling would not occur, trapping myofibre dystrophin expression in an immature state and impairing therapeutic restoration. MicroRNA candidates for regulating low-demand mRNA turnover as suggested here should instead be low abundance during early regeneration (and perhaps persistently low in dystrophic tissue), increasing only with maturity. Some studies have examined microRNA expression and contributions during early stages of muscle regeneration [61, 62], but expression data for later stages of the repair process remains limited at present.

Finally, how do myofibres sense dystrophin levels? Dystrophin protein itself is an obvious candidate: beyond physically linking the actin cytoskeleton to the ECM, sarcolemmal dystrophin plays multiple signalling roles. Dp427m mediates nitric oxide signalling via direct interaction with nNOS, and as a component of the DAGC has been linked to ion channels, G-protein coupled receptors (GPCRs), tyrosine and serine/threonine kinases, aquaporins, and other signalling proteins [63]. Sarcolemmal dystrophin could thus elicit maturity-associated pro-degradation/anti-translation signalling via one or more of these (as discussed above, similar mechanisms could suppress miR-31 expression). Signalling in this manner would be effective in a global context, i.e. during muscle regeneration or embryonic myogenesis, but might fare more poorly in local scenarios, such as stabilisation of dystrophin mRNA to produce protein for smaller repairs (discussed above). Here negative signals from adjacent ‘dystrophin sufficient’ regions might overwhelm the focal absence of such signalling from a ‘dystrophin deficient’ microtear. Signalling coupled to the absence of sarcolemmal dystrophin would conversely actively promote translation over both global and local scales, but is less mechanistically intuitive. Calcium signalling could play a role: microtears are known to cause transient but dramatic Ca^2+^ influxes [64], and defects in calcium handling are common in myopathic disorders [65-67] (indeed, aberrant sarcolemmal calcium alone can elicit Duchenne-like pathology [68]). Ca^2+^ influxes promote signalling via protein kinase C and the Rho GTPase CDC42 [41], which influence transcriptional programs in local myonuclei: concomitant modulation of post-transcriptional programs is plausible. There is moreover no reason why regulation of stability should be restricted to a single mechanism: pro-degradation signalling via correctly localised dystrophin and pro-stability signalling via calcium or other microtear marker would synergistically produce a highly responsive and flexible post-transcriptional dystrophin regulatory network.

The elaborate regulatory mechanisms proposed above all stem from the size of the dystrophin locus, raising the question: why does the gene remain so enormous? Coding sequence accounts for only ∼0.5% of the *Dmd* gene: the remainder is intronic, and thus potentially dispensable. The cost of continuous (and in most cases, superfluous) mRNA production is modest and demonstrably affordable, yet it is still a cost, and one which might be expected to drive selection pressure toward a shorter, more conventionally controllable gene. The *Dmd* gene has multiple exonic splicing enhancers and silencers (ESE/ESS) [69], thus likely carries a similar suite of intronic splicing modulators (ISE/ISS), and some introns mediate long range interactions (such as between the dp427m promoter and intron 34 [4]), but the bulk of these features are only necessary because the locus is so long. Dystrophin is phylogenetically ancient: the domain structure of the protein is well conserved throughout metazoa [15], and the C-terminal region, approximately corresponding to dp71, shares homology with the similarly conserved dystrobrevins (suggesting it is ancestral to both, and that dp427 arose as a novel fusion of a dp71-like ancestor with a long spectrin-like gene [70]). Gene length, however, is far less conserved. Within the invertebrates, both *C.elegans* and *D.melanogaster* dystrophin proteins are

∼400kDa, with actin binding, spectrin repeat and C-terminal domains [71, 72], but the corresponding genes are a mere 30kb and 136kb in length, respectively: smaller even than the mammalian dp71 locus, and readily transcribed in less than an hour. Within vertebrates, the Actinopterygii (ray-finned fishes) typically carry dystrophin genes of ∼150-300kb in size (corresponding to transcription times of 1-2 hours). Within the tetrapods, however, gene lengths become larger: amphibian and bird dystrophins are ∼1.1Mb, while marsupial and placental mammal dystrophins are 2-2.3Mb [32]. Increases in size are almost exclusively due to intron expansion, but large introns also often host novel promoters (noted even in drosophila dystrophin [71]). The largest intron in mammalian dystrophin is intron 44 (∼250kb), containing the start site for the tetrapod-specific dp140 isoform [15]. Of the remaining large introns, one contains the vertebrate-restricted dp116 promoter (intron 55, ∼125kb), but the others are upstream (introns 2, 6 and 7) i.e. restricted to dp427 isoforms. One possibility is thus a dramatic intron 44 expansion in a tetrapod ancestor, giving rise to dp140 but also substantially increasing transcription time of all upstream isoforms (addition of 250kb would effectively double the length of the fish dystrophin genes). Selective pressure to generate sufficient dystrophin against this transcriptional lag might consequently favour overproduction, with post-transcriptional control being a subsequent refinement. Here the gene would already be too large for conventional control (we have shown that cells expressing dp140 exhibit high numbers of nascent nuclear transcripts [9, 10]): further expansion of upstream introns would thus be tolerated. Conversely, reductions in gene length might result in constitutive excess of a low-demand transcript, potentially saturating degradation and translation machinery. Supportive of this, bird lineages are notorious for genomic reduction, with the pressure for lower body mass driving heavy pruning of superfluous genome sequence [73], and while bird dystrophins are smaller, they are still large. The ∼1.1Mb of avian dystrophin (∼8-hour transcription) might represent the minimum gene length that still permits the transcriptional model proposed here.

An alternative hypothesis is ‘compromise driven by necessity’: while dystrophin has multiple promoters generating multiple isoforms, all (except the 3’ truncated dp40) share the same 3’ UTR. As discussed above, 3’ UTR sequence influences stability, targeting and translational accessibility: for dystrophin, all regulatory elements within this long UTR must necessarily influence all isoforms. Inherently low mRNA stability is readily compatible with the swiftly produced dp71, allowing dynamic changes both up and down, but when applied to the lengthy dp427 transcription time, only continuous oversupply can provide the same responsiveness. Since promoter activity can conversely influence transcription in an isoform-specific manner, selective pressure for sufficient transcription to meet established post-transcriptional control systems might underpin the model presented here (notably, the brain full-length isoform dp427c also exhibits marked 5’:3’ imbalance [9, 10]).

A corollary of the above hypotheses is that we would expect our transcriptional model to apply to all tetrapod dystrophin expression, but not necessarily to the ray-finned fish, or to arthropods. Zebrafish dystrophin expression differs substantially from mammals [74], without prominent nuclear nascent transcript numbers [75]: future work assessing 5’:3’ ratios in these lineages might confirm these findings. Comparative analysis of 3’ UTR elements would also be informative: elements critical for the post-transcriptional control explored here should be conversed only between tetrapod lineages.

## Conclusion

The size of the dystrophin gene renders expression perennially problematic, with conventional regulation taking too long to deliver mRNA at times of need, yet also risking marked oversupply once demands are met: either never enough, or always too much. The work here builds toward a picture whereby myonuclei pay the 16-hour price both in advance and in perpetuity, solving the problem instead, at almost every stage, post-transcriptionally. Expression begins early to avoid delay, but translation is simultaneously suppressed (via microRNA interaction) to prevent aberrant protein accumulation prior to myotube formation. Subsequent high sarcolemmal demands are addressed by relieving translational suppression and enhancing transcript stability, ensuring large numbers of translation-competent mRNAs. Once myofibres reach maturity and demand falls, supply remains high but turnover increases to keep transcript levels low while retaining capacity for rapid responses should they be needed in future (figure 8D).

Finally, while we studied healthy tissue here, these mechanisms will also be present in dystrophic muscle. The asynchronous nature of dystrophic regeneration presumably hampers stage-specific control, as does dysregulation due to persistent dystrophin deficiency (shown for miR-31 [58]), but despite these factors, dystrophic myonuclei demonstrably continue to produce dp427 mRNA. This raises therapeutic implications, given that achieving high levels of corrected or recombinant dystrophin

mRNA is a major challenge to current approaches. Relieving early translational suppression enhances dystrophin protein levels following exon skipping [56], and our work here implies additional regulatory systems exist, all of which might be therapeutic targets. Furthermore, the extent to which truncated, therapeutic mini- or microdystrophins (which by necessity lack regulatory UTR sequences) are subject to similar controls is worthy of future study. In summary though, high dystrophin demand appears to be primarily met by maximising translational value of transcripts, regardless of numbers: pharmacological modulation of mRNA turnover and translation might be a means to achieve beneficial protein restoration with even low efficiency correction at the mRNA level.

## Author contributions

JCWH: Conceptualization, Formal Analysis, Funding Acquisition, Investigation, Visualization, Writing - Original Draft Preparation; LER: Formal Analysis, Investigation, Visualization, Writing – Review & Editing; DJW: Resources, Supervision, Writing – Review & Editing; RJP: Funding Acquisition, Supervision, Writing - Review & Editing

## Supplementary figure legends

Supplementary figure 1: Dystrophin protein remains absent at 4 DPI

At 4 DPI, fibre profiles can be identified via perlecan staining, but sarcolemmal dystrophin staining is essentially absent from all damaged fibres (which are concomitantly robustly positive for infiltrating IgG). Scalebars: 100µm

Supplementary figure 2: Dystrophin ISH signal is absent at 2 DPI

Multiplex labelling of dystrophin mRNA (5’: green; middle: yellow; 3’: magenta) in injured muscle at 2 days post injury reveals no labelling with any probe, indicating absence of both mature and nascent transcripts.

Supplementary figure 3: *Pax7* expression is not detected at 2 DPI

Multiplex labelling of dystrophin and *Pax7* mRNA (dp427 5’: green; dp427 3’: magenta; *pax7*: yellow) in injured muscle at 2 days post injury reveals no labelling with any probe, indicating absence of nascent and mature dp427 transcripts, and no detectable *pax7* expression.

Supplementary figure 4: *Ki67* expression is minimal in healthy muscle

Multiplex labelling of dystrophin and *Ki67* mRNA (dp427 5’: green; dp427 3’: magenta; *ki67*: yellow) reveals the characteristic strong nuclear 5’ signals of nascent dp427 mRNAs and the punctate sarcoplasmic 5’ and 3’ signals of mature transcripts, while Ki67 labelling is very low and highly sporadic, found only within rare individual nuclei.

