## Supplementary figures and images for "Spatiotemporal analysis of dystrophin expression during muscle repair"

### Supplementary Figure 1

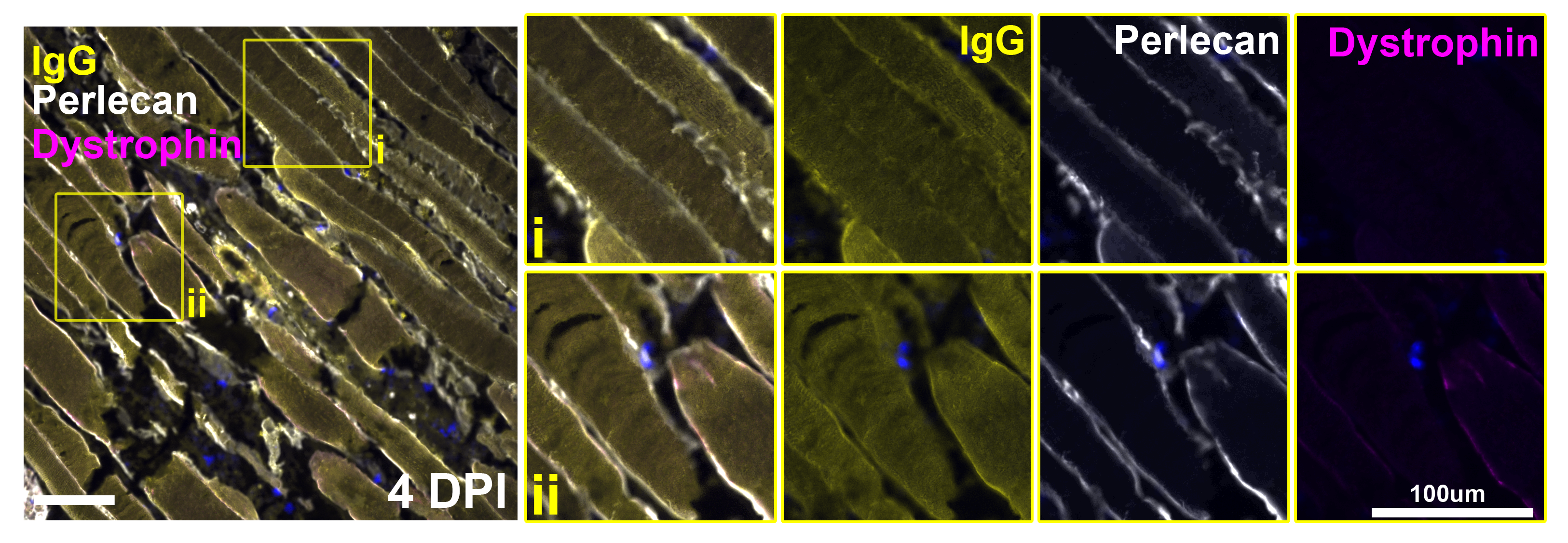

### Supplementary Figure 2

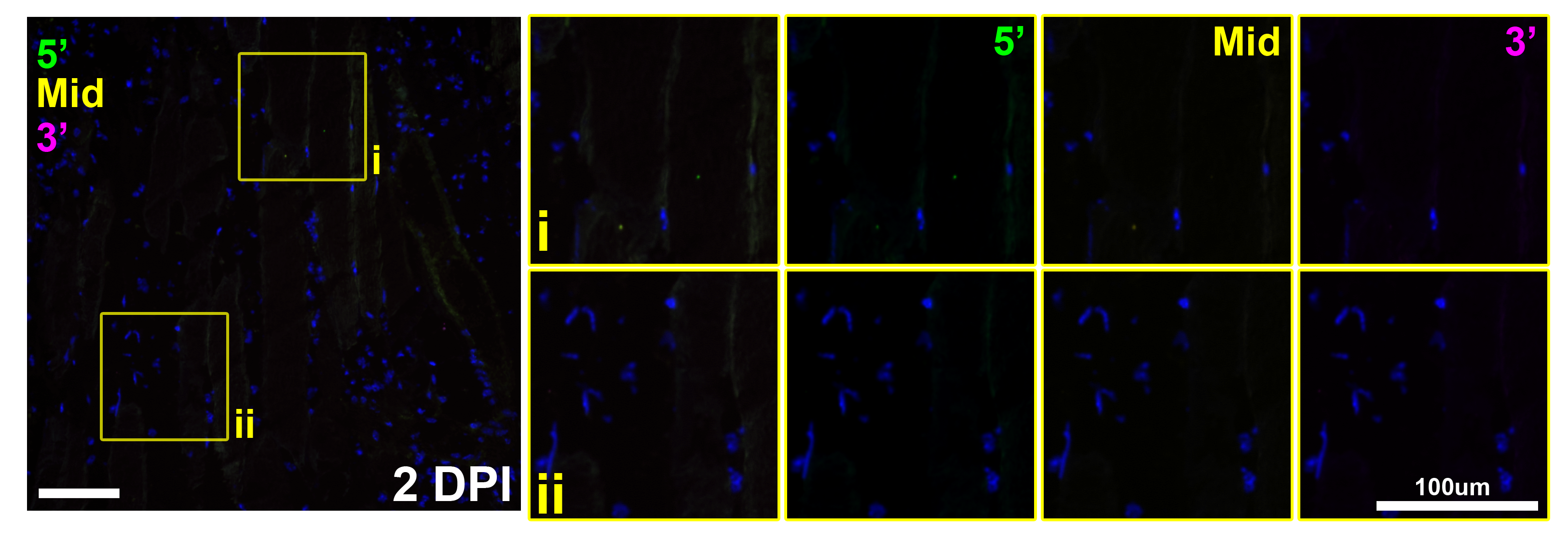

### Supplementary Figure 3

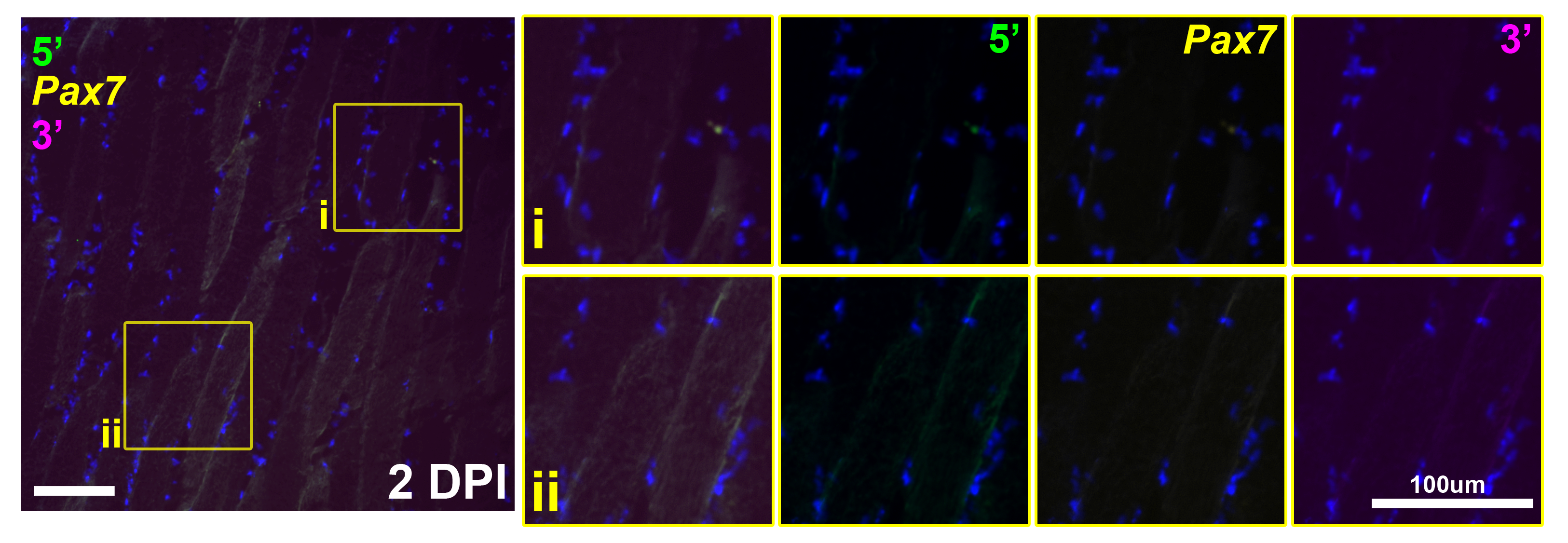

### Supplementary Figure 4

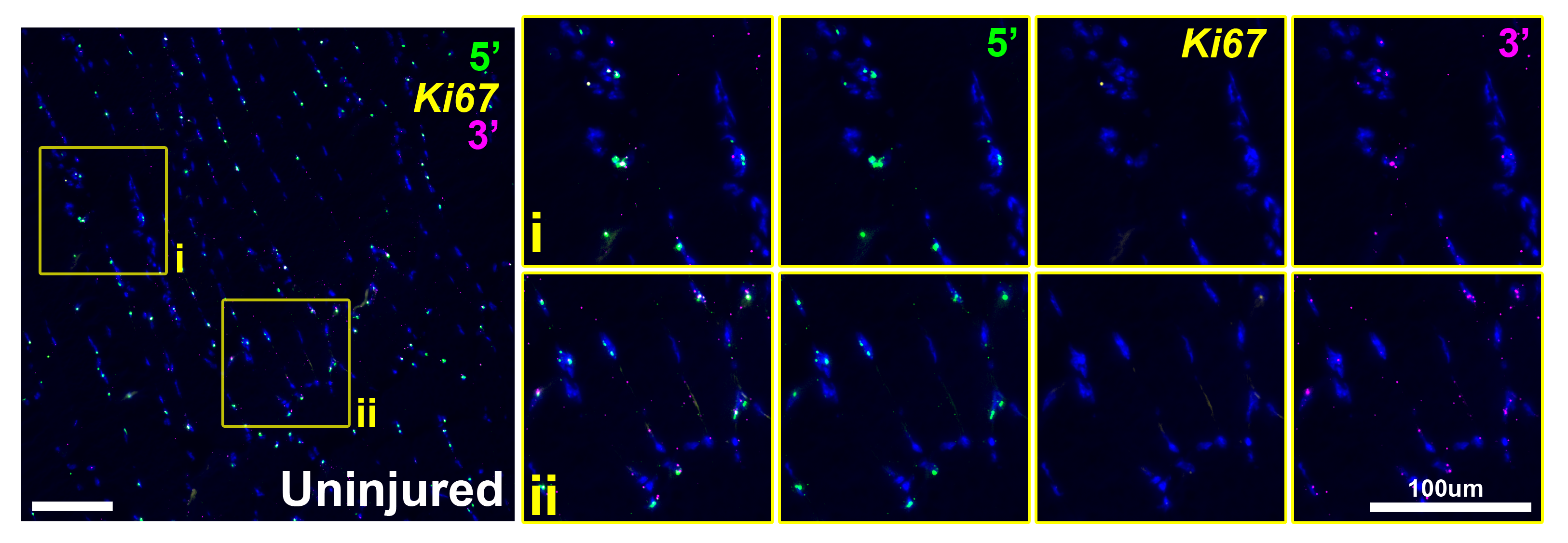
